# Resolving the haplotypes of arbuscular mycorrhizal fungi highlights the role of two nuclear populations in host interactions

**DOI:** 10.1101/2023.01.15.524138

**Authors:** Jana Sperschneider, Gokalp Yildirir, Yanina Rizzi, Mathu Malar C, Essam Sorwar, Eric CH Chen, Wataru Iwasaki, Elizabeth K. Brauer, Whynn Bosnich, Caroline Gutjahr, Nicolas Corradi

**Author notes:** Max-Planck-Institute of Molecular Plant Physiology, Am Mühlenberg 1, 14476 Potsdam-Golm, Germany (Yanina Rizzi, Caroline Gutjahr). Contributed equally (Jana Sperschneider, Gokalp Yildirir).

## Abstract

Arbuscular mycorrhizal fungi (AMF) are prominent root symbionts with a multinucleate cytoplasm that can carry thousands of nuclei deriving from two parental strains and varying in relative abundance in a large syncytium. Here, we set out to improve our understanding of such remarkable genetics by resolving the nuclear genomes of all publicly available AMF heterokaryons using PacBio HiFi and Hi-C sequencing. We find that all AMF heterokaryons carry two sets of homologous chromosomes, where genes associated with plant colonization reside in gene-sparse, repeat-rich compartments. The co-existing nuclear genomes are phylogenetically related but differ significantly in content and epigenetics, resulting in nucleus-specific regulatory programs during mycorrhizal interactions. AMF heterokaryons carry signatures of past genetic exchange indicative of sexual reproduction, followed by clonal haplotype evolution. This work uncovers the contribution and origin of nuclear genomes present in AMF heterokaryons and opens avenues for improvement and environmental application of these strains.

## Introduction

Arbuscular mycorrhizal fungi are prominent root symbionts of the subphylum Glomeromycotina ^1^ that can improve access to nutrients and resistance against pathogens in land plants ^2^. These keystone mutualists increase ecosystem productivity in global terrestrial ecosystems ^3^ and many industries specialize in their production as bio-stimulants for agricultural practices, forestry, and plant nurseries. AMF can contain over 20,000 nuclei within individual spores and, as opposed to other multinucleate fungi, there is no stage in their life cycle where spores and cells contain one or two nuclei ^4^. Fossil and phylogenomic evidence indicates this plant-fungus symbiosis dates back more than 400 million years ^5,6^, but despite this longevity no sexual apparatus has been observed in these organisms. Because of this AMF were long pigeonholed in a selected group of organisms called “ancient asexuals” ^7^, and this absence of sexual reproduction was proposed to be offset by high degree of sequence divergence among their co-existing nuclei ^8^. The view that AMF have a purely clonal lifestyle and non-mendelian genetics has, however, been challenged by discoveries of gene orthologues that function in meiosis ^9^, and more recently by evidence that some strains of the model species *Rhizophagus irregularis*, referred to as AMF heterokaryons (or AMF dikaryons) carry thousands of nuclei deriving from two parental strains ^10^. This genetic state is analogous to the heterokaryotic (sexual) stage of the Dikarya (ascomycete and basidiomycete fungi) where two or more parental nuclei co-exist in one cell for multiple generations ^11^, and indeed AMF heterokaryons share remarkable genomic features, ecological traits and nuclear dynamics with heterokaryotic fungal relatives ^4^.

For example, the parental nuclei of fungal heterokaryons carry diverging mating-type loci (MAT-loci) that orchestrate sexual interactions between compatible strains, and loci with remarkable similarity are also present in nuclei with distinct genotypes in AMF heterokaryons ^10,12^. The co-existence of two nuclear types in fungal heterokaryons improves fitness and adaptability to environmental change, and this ability correlates with change in the relative ratio of co-existing nuclei within the mycelium ^13,14^. Consistent with this, AMF heterokaryons outperform homokaryotic relatives in many life-history traits ^15^, including hyphal growth speed and density, and their two haplotypes also vary in relative abundance depending on environmental factors and host identity ^16,17^.

In the dikaryotic basidiomycete *Agaricus bisporus* and the heterokaryons of *Neurospora tetrasperma*, two parental nuclei express different genes throughout the life cycle ^18,19^, and in other species allele-specific expression is tightly linked to the ratio of the two nuclear types ^20^. These findings show that parental genomes play distinct molecular roles in the biology and nuclear dynamics of fungal heterokaryons, but whether similar mechanisms also exist in AMF heterokaryons is unknown because sequence information on parental nuclei is unavailable. As a result, their relative genetic and epigenetic contribution to the biology of these strains and their mycorrhizal partners could thus far not be investigated.

To address these questions, we resolved the parental nuclear sequences of all publicly available AMF heterokaryons using PacBio High Fidelity (HiFi) reads and Hi-C sequencing and analyzed their gene regulation across multiple hosts and conditions. Untangling each of the haploid genomes also allowed determination of the evolutionary origin of these strains, and examination of whether AMF heterokaryons carry two dominant parental genomes that physically separate across thousands of nuclei, as proposed by Ropars and colleagues ^10^.

## Results

### Nuclear genome phasing and annotation of AMF heterokaryons

HiFi PacBio long-read and Hi-C sequencing data was obtained for all publicly available AMF heterokaryons (A4, DAOM-664343; A5, DAOM-664344; G1, DAOM-970895; SL1, DAOM-240409). This data was assembled with hifiasm in Hi-C integration mode ^21^, which returned two haplotypes for each strain. Phasing was assessed with NuclearPhaser ^22^ and after minor phase switching correction the assembly was scaffolded into 32 chromosome-scale scaffolds, guided by Hi-C data (**Table 1, Figure 1, Supplemental Figures 1-4)**.

**Table 1:**
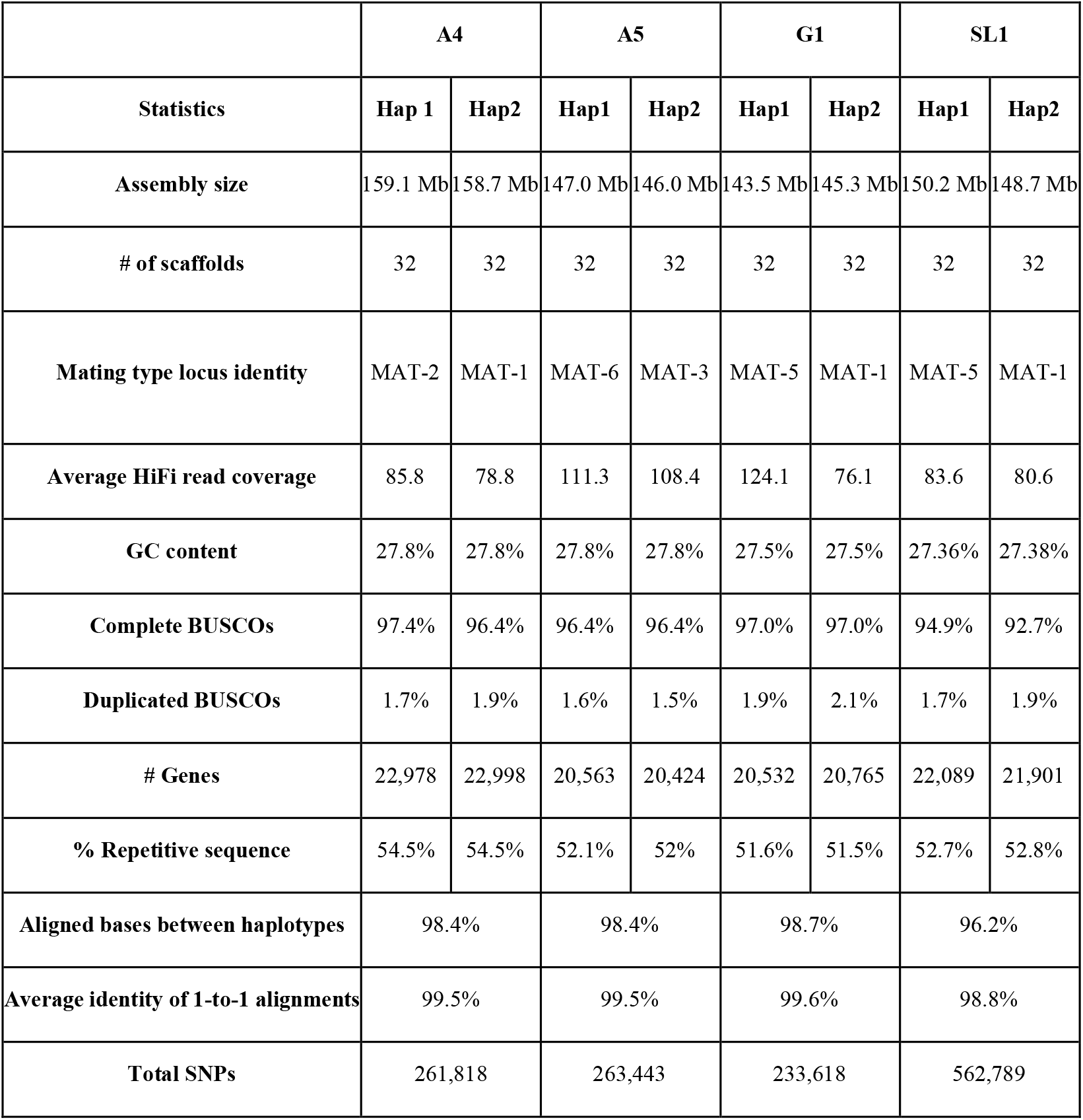
General genome assembly statistics for the AMF heterokaryons.

**Figure 1:**
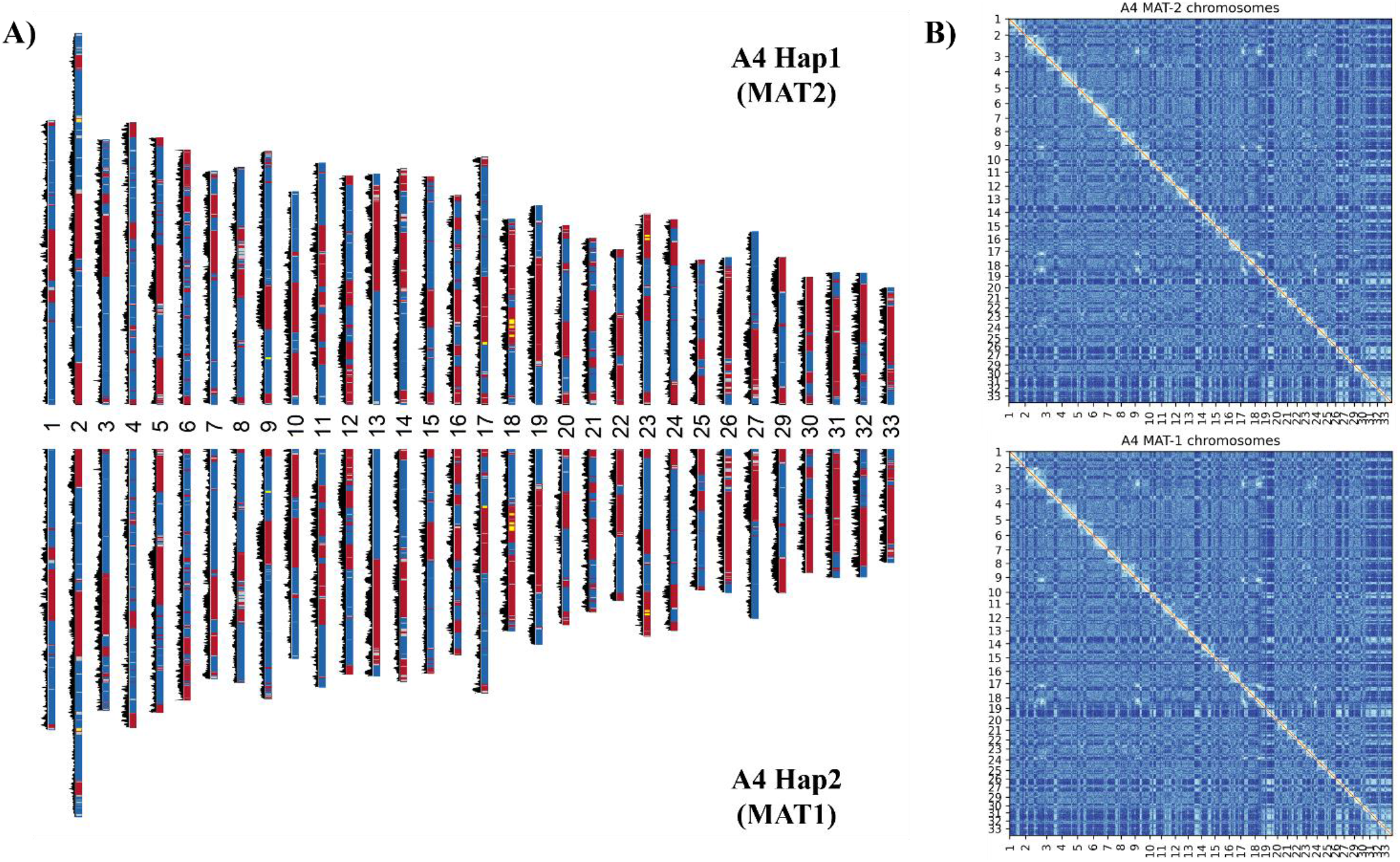
The phased chromosomes of the AMF heterokaryons. **(A) The chromosomes of the A4 haplotypes.** Gene density is shown in black (20 Kb windows). A and B compartments are shown in red and blue, respectively. rRNA operons are shown in yellow. All haplotypes carry 32 chromosomes and A/B compartments. **(B) The Hi-C contact map of the A4 haplotypes shows a compartmentalization of the chromosomes into A and B compartments.** A ‘checkered’ pattern, resulting from the interactions of several chromosomes, indicates the genome arrangement of the euchromatin or heterochromatin compartments of *R. irregularis*. Data from other isolates is available in Supplemental Figures 2-4.

On average, only 54 short contigs totaling 2.7 Mb (L50: 61 Kb) remained un-scaffolded per strain. Consistent with phasing of fungal dikaryons based on Hi-C data ^22–24^, the haplotypes of AMF heterokaryons exhibit a strong dikaryotic phasing signal, with ~90% of inter-chromosome (*trans*) Hi-C contacts contained within nuclear haplotypes (**Supplemental Figures 1-4)**. In support of molecular analyses of single spores ^17^, the MAT-2 and MAT-5 haplotypes have higher coverage (relative abundance) in the isolates A4 and G1, respectively (~9.2% and 63% higher) but are closer to balance in A5 and SL1 in extraradical mycelium after propagation with *Daucus carota cv* P68 (Carrot; 3.5% average difference) (**Table 1**). Analysis of allele frequencies confirm these findings (**Supplemental Figure 5)** and showed that SNPs with frequencies diverging from the expected two haplotypes - i.e. putative somatic diversity affecting individual nuclei - are exceedingly rare (average 0.005% of the non-repeated portion of the genome) ^10,12^.

In our assemblies, one part of the chromosome 28 previously identified in AMF homokaryons ^25^ maps to chromosome 2 and the other to chromosome 17, but inspection of Hi-C contact maps and HiFi read coverage did not provide conclusive evidence that these chromosomes should be broken. The identification of AMF centromeres will ultimately be necessary to determine if *Rhizophagus irregularis* carries 32 or 33 chromosomes ^26^, and for ease of comparative analysis we numbered the chromosomes according to the homologous chromosomes of AMF homokaryons ^25^

Phased haplotypes have genome sizes and number of genes and repeats in expected range for the species and their complete BUSCO gene repertoire averages 96% ^10^ (**Table 1**). Unlike other fungi, AMF do not carry thousands of homogenous ribosomal operons (rRNA), but harbor up to 11 divergent rRNA copies on each haplotype ^25^ (**Figure 1, Supplemental Table 1**). The Hi-C contact maps confirm that chromosomes of AMF heterokaryons separate into two dominant euchromatin (A) and heterochromatin (B) compartments known to dictate the genome biology and evolution of these symbionts ^25^. The A-compartment has higher gene density, while the B-compartment has higher repeat density and is enriched in secreted proteins and candidate effectors, including mycFold effectors ^27^ (**Figure 1, Supplemental Figures 2-4, Supplemental Table 2, Supplemental Table 3**).

**Figure 2.**
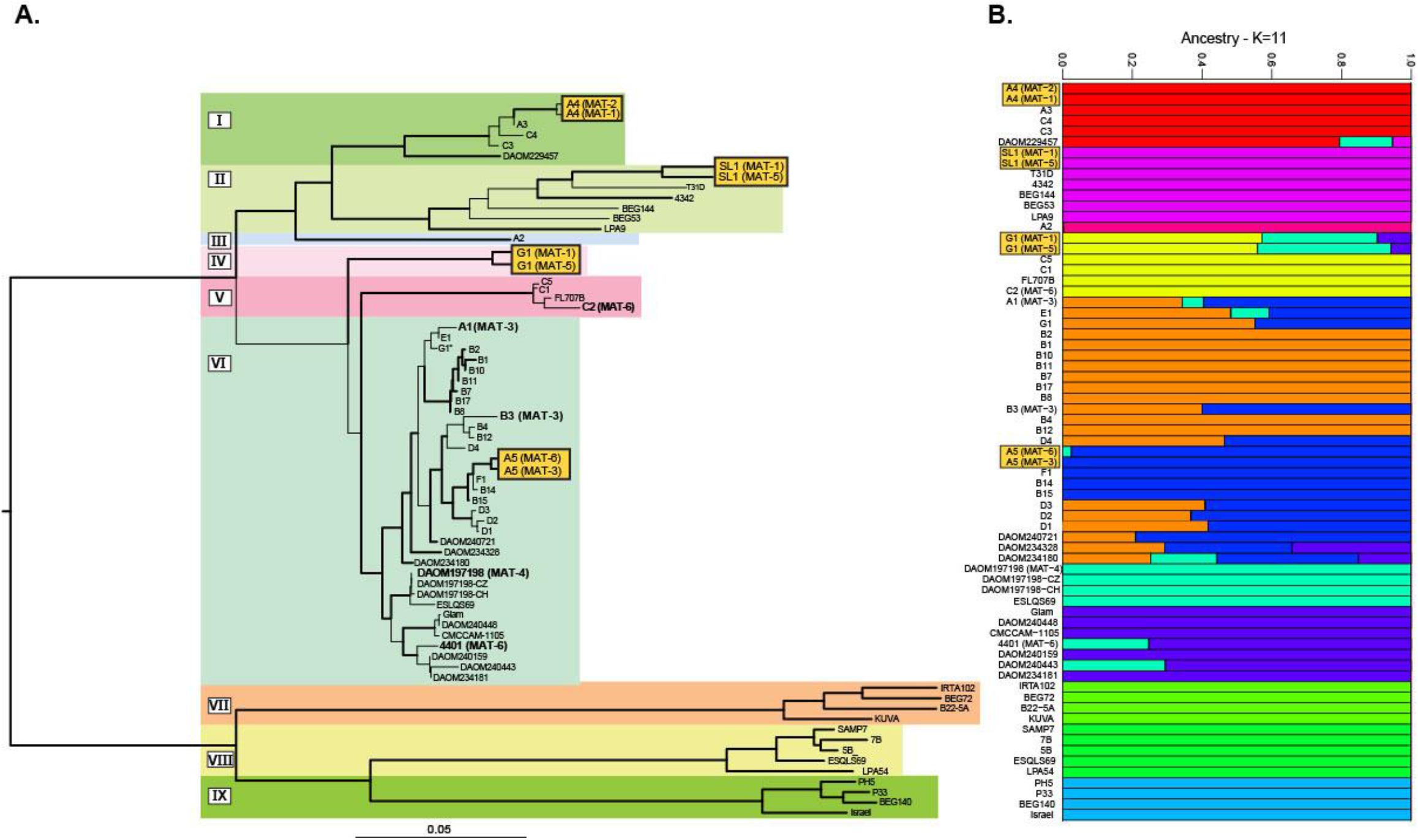
Phylogenetic tree constructed with 65 *R. irregularis* strains (incl. 8 haplotypes from AMF heterokaryons). Haplotypes from AMF heterokaryons are shown in yellow squares. Data obtained using chromosome-level genome assemblies are shown in bold. Based on relative branch lengths, the phylogeny resolves 9 clades which are highlighted in color. Thicker branches indicate bootstrap support > 95. The tree was made using IQTREE algorithm, in GTR-FO mode with 1000 bootstrap replicates. Scale bar represents 0.04 substitutions per site. The G1 strain located in clade VI and noted with an asterisk is homokaryotic and does not represent the heterokaryotic isolate G1 (DAOM-970895) from clade IV. **B. Population structure of R. irregularis at K = 11.** The structure has been inferred using NGSadmix. Each row represents a strain, listed as in the ML tree (A). Colored bars represent their coefficients of membership in the various gene pools based on SNP data.

### Phylogenetic and population analyses of AMF heterokaryons

The phased haplotypes were first used to assess the evolutionary origin of AMF heterokaryons. A maximum likelihood phylogeny was generated using available genome and RAD-seq data from 65 strains catalogued as *R. irregularis* ^25,28^ (**Figure 2**), which revealed at least 9 supported phylogenetic clades for this species.

Consistent with past analyses, *R. irregularis* strains do not always cluster in the phylogeny according to geographical location or mating-type identity ^28,29^. All AMF heterokaryons cluster into separate clades, but their co-existing haplotypes always group together and away from other sequenced strains, meaning that their parental homokaryotic strains could not be identified with available datasets. Long branch lengths indicative of long-term clonal divergence separate some co-existing haplotypes, especially G1 and SL1. Inspection of aligned homologous positions used to produce the phylogeny did not reveal SNP tracts shared between co-existing haplotypes and other strains that could indicate the presence of recombination following the formation of AMF heterokaryons. However, evidence of admixture indicative of past genetic exchanges (e.g., recombination, introgression) exists between multiple strains. In AMF heterokaryons, evidence of past admixture events is found in both haplotypes of G1, and a small portion of the MAT-6 haplotype in A5.

### Comparative analyses of co-existing haplotypes

Large-scale chromosome reassortments exist between co-existing haplotypes, and at smaller genome scale variability in gene order and content is common. While homologous chromosomes can experience inversions and translocations (**Figure 3a, Supplemental Figure 6**), at the putative AMF mating type locus only five genes (a hypothetical protein, a choline transporter, the homeodomains HD-1 and HD2, and a phosphoglycerate mutase) are colinear among all haplotypes (**Figure 3b**).

**Figure 3.**
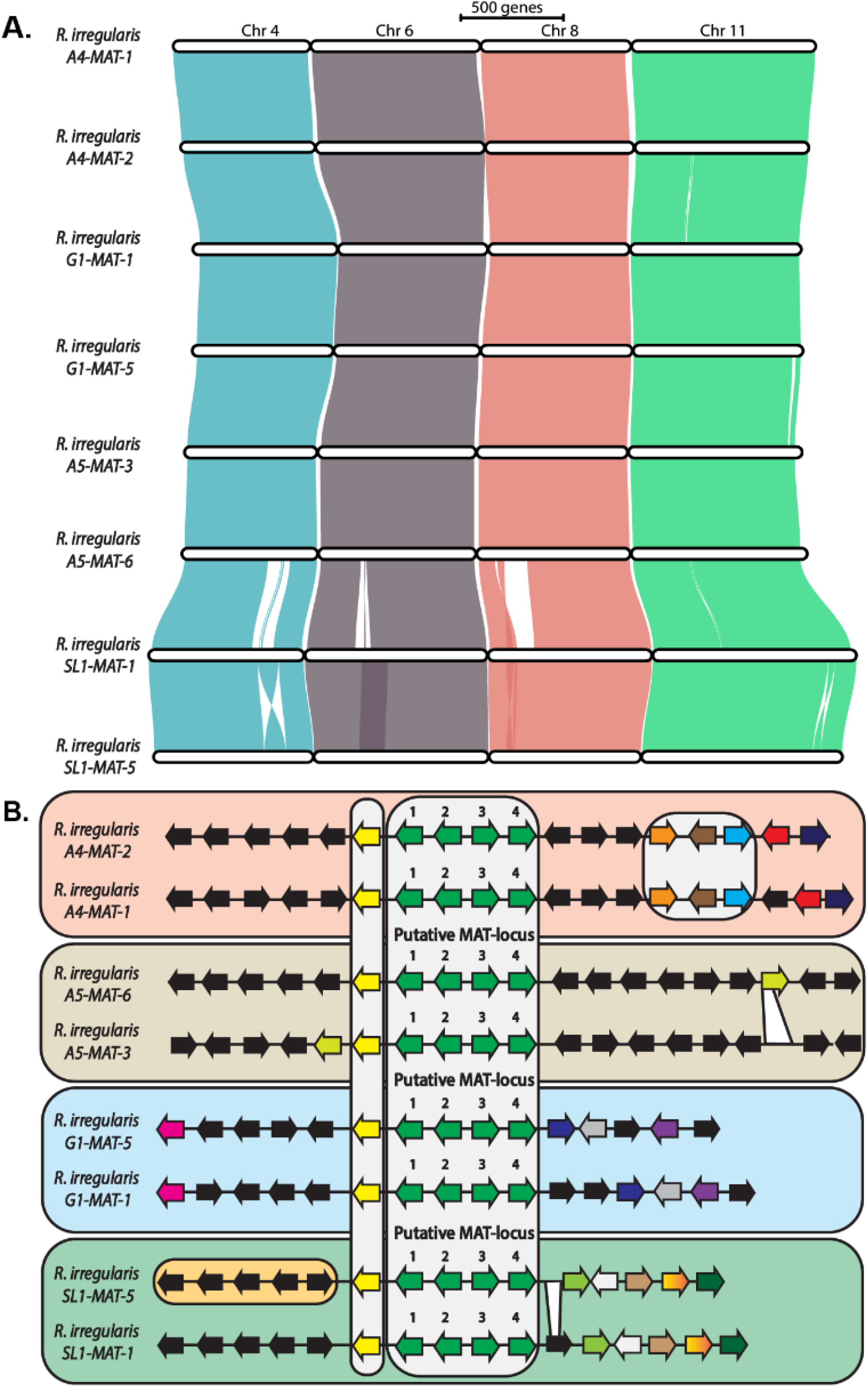
A. Gene synteny and reassortments in representative chromosomes (4, 6, 8 and 11) of AMF heterokaryons. Similar patterns affect all chromosomes **. B. Gene order is relaxed in regions surrounding the putative MAT-locus of all AMF heterokaryons.** The four genes thought to compose the AMF MAT-locus (located on chromosome 11) are shown as green arrows and highlighted by numbers as follows; 1. Choline transporter, 2. HD1, 3. HD2, 4. phosphoglycerate mutase. Arrows indicate gene transcriptional direction and color highlights sequence orthology among haplotypes. Black arrows indicate genes showing no evidence of orthology among co-existing haplotypes. SL1-MAT-5 orthologs within orange ellipse are located 20Kb upstream in the MAT-1 haplotype.

AMF heterokaryon haplotypes also differ in relative gene function and number of repeats. For instance, in the strain A4 we observed an enrichment of the tetratricopeptide repeat domain Sel1 on MAT-2, whereas an enrichment of protein kinase domains is found on MAT-1 **(Supplemental Figure 7, Supplemental Figure 8)**. All strains also carry genes specific to each haplotype. These include candidate effectors and secreted proteins potentially involved in the dialogue between the mycorrhizal partners, as well as rRNA operons and repeats.

In the strain G1, the MAT-5 haplotype has one more rRNA operon, but 233 less genes than the MAT-1 haplotype (**Table 1, Supplemental Table 1**). Moreover, co-existing haplotypes can differ in chromosomal compartmentalization (**Supplemental Figure 9).** Despite these differences, haplotypes are more similar in content and chromosome size than different *R. irregularis* strains with available genomes are to one another ^25^. For instance, co-existing haplotypes differ by an average of 145 genes and 578 kilobases (Kb) of repeats in AMF heterokaryons compared to 2,646 genes and 16.4 megabases (Mb) of repeats between AMF homokaryons, and their homologous chromosomes differ in size by just over 100 Kb compared to over 550 Kb for AMF homokaryons ^25^.

### Haplotype-specific gene expression and regulation across hosts

We hypothesized that the observed genetic and epigenetic differences affect the transcriptome of AMF heterokaryons. To assess this, we obtained RNA-sequencing data for all AMF heterokaryons in the following conditions: extraradical mycelium isolated from root organ cultures (ROC), colonized ROC of *D. carota* cv. P68 (carrot) and *C. intybus* (chicory), and colonized roots of whole plants from the dicotyledon *Lotus japonicus* (Lotus) and the monocotyledon *Brachypodium distachyon* (Brachy). We first assessed the localization of differentially expressed genes on the chromosomes, i.e. the A/B compartments ^25^. Over 40% of the genes encoding secreted proteins up-regulated during *in planta* colonization, including all but one mycFOLD effector, are preferentially located in the repeat-rich B-compartment but also preferentially on one of the haplotypes (**Supplemental Figure 10**).

To determine the relative contribution of each haplotype, we used approaches that conservatively measure the expression of orthogroups with exactly one gene copy on each haplotype and where the orthologs can be differentiated by at least one SNP in their coding sequences ^18,19^ (A4: 8,064 genes; A5: 7,152 genes; G1: 6,816 genes). Average expression and regulation of these 1-1 orthologs can differ significantly among co-existing haplotypes across hosts and conditions (**Figure 4ab, Supplemental Figure 11**). Overall, total and average gene expression are consistent with the relative haplotype abundance measured by relative coverage (**Table 1, Supplemental Figure 5**) and digital droplet PCR (ddPCR) on the same colonized roots (**Supplemental Figure 12**), that is, the most abundant haplotype has higher total expression. Specifically, in the strains A5, A4, and G1, 1-1 orthologs from the MAT-3, MAT-2 and MAT-5 haplotypes have higher average gene expression across all conditions (not significant with chicory for A5 and A4). Identical patterns are seen for A5, SL1 and G1 with whole plants, indicating that similar nuclear dynamics and relative expression occur in between colonized roots of ROC and whole plants for these strains (**Figure 4a, Supplemental Figure 11)**.

**Figure 4:**
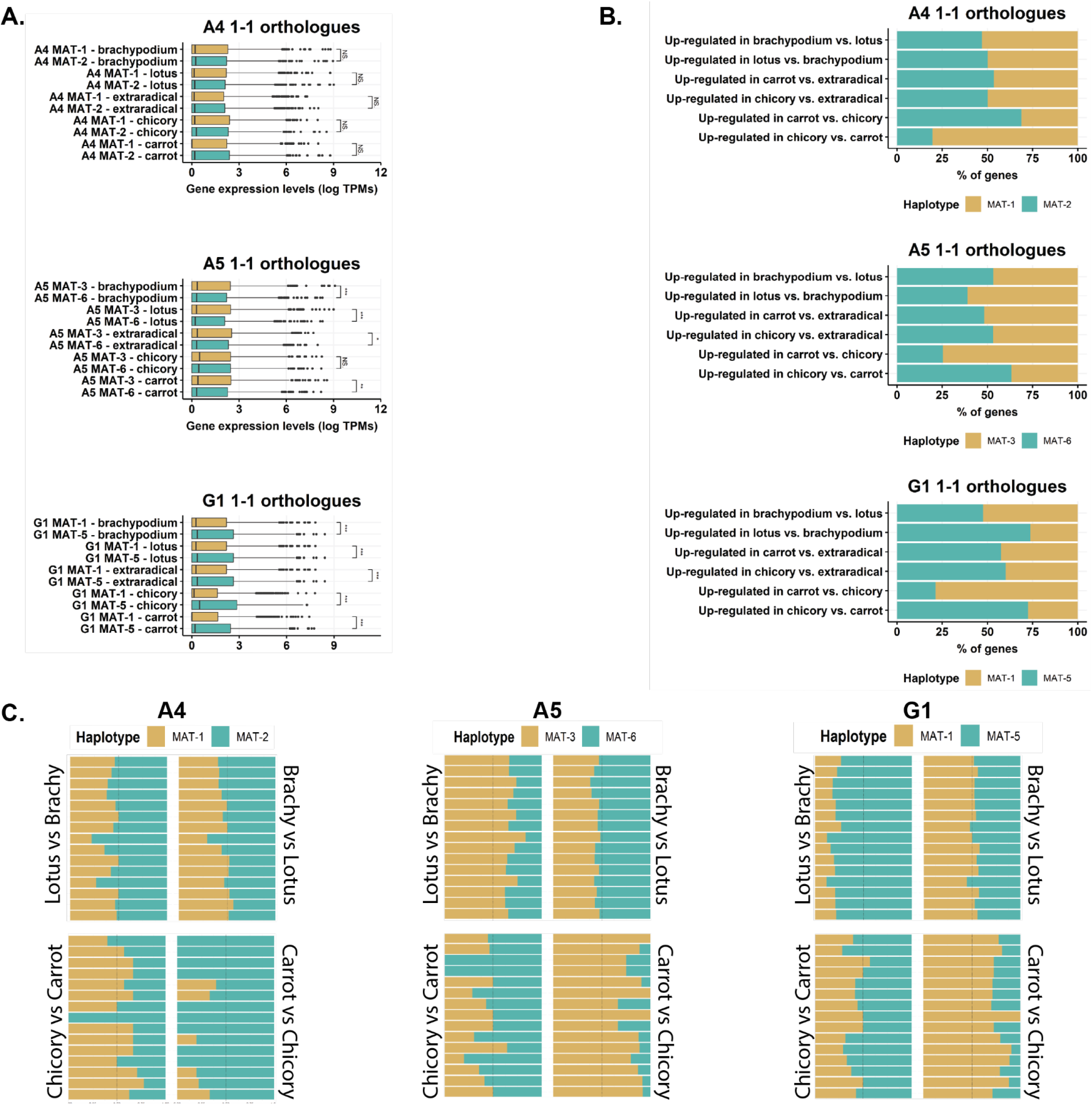
Haplotype specific RNA-seq analyses for A4, A5 and G1. A. Haplotype-specific expression. (left) of 1-1 orthologues (right). Box plot edges show the first and third quartile in box plots, with the median shown as the middle lane. The data range is indicated by whiskers and dots represent outliers. Asterisks indicate a t-test statistical significance of at least P < 0.05**. B. The % of genes upregulated in each haplotype** across the three conditions used in this study (in planta-carrot, in planta-chicory and extraradical mycelium). Similar analyses for SL1 are available as supplemental figure 11**. C. GoTerm enrichment comparisons of 1-1 orthologues** between haplotype specific genes upregulated in carrot versus chicory (upper box) and chicory vs carrot (bottom box) in A4, A5 and G1. Each bar represents percentage of upregulated genes for the same GoTerm in each haplotype. Only GoTerms shared among all haplotype-specific upregulated genes are shown to highlight the respective functional contribution of each haplotype across hosts. List of shared GoTerms presented in Figure 4C is available as Supplemental Table 4. Enrichments of GoTerms unique to each haplotype also exist (Supplemental Figure 13).

With few exceptions, haplotype-specific gene regulation follows this trend based on host identity (**Figure 4b, Supplemental Figure 11**). For example, in A4, 68.9% of the 1-1 orthologs up-regulated in carrot compared to chicory are located on the more abundant MAT-2 haplotype, but 80.3% of the 1-1 orthologs that are up-regulated in chicory compared to carrot are located on the MAT-1 haplotype which has slightly lower abundance in colonized roots. In A5, 63.4% of the 1-1 orthologs that are up-regulated in chicory compared to carrot are located on the slightly more abundant MAT-6 haplotype, and again the inverse pattern is seen for those up-regulated in carrot compared to chicory (74.6% on the more abundant MAT-3 haplotype).

Remarkably, in G1 78.6% of 1-1 orthologs on the MAT-1 haplotype are up-regulated with carrot compared to chicory despite this strain having its nuclei belonging overwhelmingly to MAT-5 (**Figure 4b, Table 1, Supplementary Table 12**). Similar patterns exist in whole plant for all strains. In A5, 53.5% of 1-1 orthologs up-regulated in brachy compared to lotus are on the MAT-6 haplotype, but 61% of those are on MAT-3 haplotype in lotus compared to brachy (**Figure 4b, Supplemental Figure 11**). In G1, 52.3% of 1-1 orthologs up-regulated in brachy compared to lotus are on the MAT-1 haplotype, but 73.8% of those are on MAT-5 in lotus compared to brachy (**Figure 4b**), and in SL1 1-1 orthologs located on the MAT-1 and MAT-5 haplotypes are equally up-regulated in brachy compared to lotus, but over 65% of those are on the MAT-5 haplotype in lotus compared to brachy (**Supplemental Figure 11**).

The observed variability leads to distinct putative functional contributions for co-existing haplotypes across growth conditions (**Figure 4c**). For example, in the strain A4, the genes upregulated in the MAT-1 haplotype in *C. intybus* compared to *D. carota* are enriched for GO-terms involved in reproduction, cellular process and cellular component organization, but not in *D. carota* as those are primarily upregulated in the MAT-2 haplotype (**Figure 4c)**. The same patterns are seen in A5, where genes with putative functions in reproduction and developmental process are upregulated primarily by genes located in the MAT-3 haplotype in *D. carota* and the MAT-6 haplotype in *C. intybus*. Similarly, regulation of GO-terms is enriched in the MAT-3 haplotype with *L. japonicus* in A5, but is more balanced among haplotypes in *B. distachyon.* In G1, upregulated GO-terms are overwhelmingly controlled by MAT-5 haplotype in *L. japonicus*, but not with *B. distachyon*. Each haplotype also regulates unique functions during root colonization and in the extra radical mycelium, comparisons against extraradical mycelium show a generally more balanced haplotype contribution to each GO-Term (**Supplemental Figure 13**).

## Discussion

The nuclear organization of AMF has been a long source of debate ^30^. The recent identification of homokaryotic and heterokaryotic strains reconciled previous views in this area ^10^, but these findings were built on highly fragmented genome and nuclear datasets that may skew data interpretation and conclusions. Here, we show that combining PacBio HiFi long reads with Hi-C sequencing phases two complete sets of chromosomes in all AMF heterokaryons. These haplotypes physically separate among thousands of individual nuclei, which confirms current models that explain AMF genetics ^10,30^. The absence of haplotype degeneration and intra-haplotype genetic variation contrasts with predictions that long-term clonal evolution in AMF should lead to extensive nuclear divergence through accumulation of deleterious mutations ^8,31,32^. What maintains haplotype integrity in the absence of formal reproduction in AMF needs further investigation, but this feature may be achieved via mutation suppression mechanisms recently reported in dikaryotic fungi ^33^.

The fragmentary and collapsed nature of genomic data from AMF heterokaryons also hampered the identification of their evolutionary origins. By untangling co-existing haplotypes, we were able to confirm that none of these strains are closely related to homokaryons with sequenced genomes ^10^, but emerged instead from phylogenetically close (and yet to be sequenced) homokaryotic relatives with distinct MAT-loci. This information now provides a clear roadmap that can be used to identify and cross suitable candidates and produce new strains tailored to plant growth.

Haplotype sequence data from dozens of *R. irregularis* strains supports evidence of past genetic exchange between individual strains of this species. These presumably arose via meiosis-like mechanisms, and similar events, combined with signatures of clonal haplotype evolution, are also found in the parental nuclei of some AMF heterokaryons. In the absence of data from parental strains, and with no signatures of recombination involving co-existing haplotypes, only assumptions can be made for the emergence of AMF heterokaryons. As extensive divergence in gene order and structure exists among haplotypes, one possibility is that these strains did not undergo meiotic recombination and emerged instead through somatic nuclear exchange; a process that generates heterokaryotic progenies that seldom recombine via sexual reproduction in basidiomycetes ^11,34^, resulting in haplotypes evolving clonally over time. Alternatively, if sequence divergence still allows recombination amongst haplotypes, then past mating between two reasonably diverged nuclei may have produced these heterokaryons, followed by some meiosis-like events and clonal haplotype evolution for some time.

Our RNA-seq analyses clarified the relative contribution of co-existing haplotypes in AMF heterokaryons. In distant fungal heterokaryons, the co-existence and regulation of two parental genotypes improves fitness and adaptability to change ^35–37^. The variability in content, expression, and epigenetics among haplotypes is thus likely to provide similar benefits to AMF heterokaryons. In support of this, changes in host identity or growth condition alter nuclear dynamics and expression in all strains, which can result in upregulation being primarily controlled by the least abundant haplotype. As such, our work indicates that co-existing haplotypes act as distinct and active regulatory units in AMF heterokaryons, and this may explain their ability to outgrow homokaryotic relatives across a wider host range ^15,38^. Future studies should now investigate relative haplotype expression with additional varying parameter (temperature, pH, salinity, hosts) to obtain a more global view of the molecular interactions that occur between haplotypes and the surrounding environment.

Overall, this study resolved long-standing questions on the evolutionary genetics and origin of AMF heterokaryons, and now opens opportunities to investigate the nuclear biology and dynamics of these peculiar strains using microscopy and haplotype-specific probes. While evidence of past genetic exchanges in many *R. irregularis* strains and haplotypes supports the view that AMF can recombine with close relatives, the acquisition of sequence data from parental homokaryotic strains will be required to determine whether AMF heterokaryons originate because of sexual or clonal processes (or both). The finding that nuclear regulation is controlled by two haplotypes further underpins similarities between the genetics of AMF and Dikarya, which recently have been proposed to be sister clades by phylogenomic analyses ^39,40^.

## Material and Methods

### Culturing, DNA extraction from mycelium and Hi-Fi and Hi-C sequencing

*Rhizophagus irregularis* strains A4, DAOM-664343; A5, DAOM-664344; G1, DAOM-970895; SL1, DAOM-240409) were cultured using either *Daucus carota* (*cv 68*) or *Cichorium intybus* root organ cultures (ROCs) as host ^17,41^. The strains A4 and A5 were originally collected from a field in Tänikon ^42^. The SL1 and G1 strains were originally collected by staff at Agriculture and Agri-Food Canada (Ottawa, Canada) in Sainte-Foy (Québec, Canada). All strains were propagated in two-compartment ROCs, allowing us to produce mycelium and spores without obvious contaminants. High-molecular-weight DNA was extracted for all strains using a protocol proposed by Schwessinger & McDonald ^43^, and over 10 micrograms of DNA was subjected to high fidelity (HiFi) PacBio sequencing at Discovery Life Science (Huntsville, AL). PacBio sequencing produced an average 24,647,320,191 base pairs (bp) of HiFi reads with read median quality of Q33 and median read length of 13,138.

To obtain high-quality Hi-C data, approximately 200 mg of fresh mycelium from each strain was crosslinked and shipped to Phase Genomics (Seattle, WA, USA) to obtain strain-specific Hi-C data. Hi-C data were produced using the Proximo Hi-C Kit (Microbe) at Phase Genomics with DPNII restriction enzymes. Before generating high-throughput Hi-C Illumina data, the quality of these libraries was assessed by mapping low-coverage paired-end data onto available *R. irregularis* assemblies. In all cases, the values obtained for our samples far exceeded the quality thresholds set by Phase Genomics.

### Cultivation of *R. irregularis* strains with *L. japonicus* and *B. distachyon*

Seed germination and inoculation with *R. irregularis* strains was performed according to Torabi et al., (2021) ^44^. *L. japonicus* seeds were manually scarified with sand paper and surface sterilized with a solution containing 0.3% NaClO and 0.1% SDS for 5 minutes and then washed five time with sterile distilled water. Imbibed seeds were germinated on Gamborg B5 medium (Duchefa Biochemie) containing 0.8% agar and 2% sucrose at 22°C for 3 days in dark and 7 days in light (16-h-light /8-h-dark).

*Brachypodium distachyon* seeds were soaked in water for 1h and then the glume was removed. Subsequently seeds were surface sterilized with a 30 sec rinse in 70% ethanol, a 3 min 0.3% NaClO and 0.1% SDS followed by five rinses with sterile distilled water. Seeds were transferred to water-saturated Whatman paper in a Petri dish, and the dish was sealed with parafilm. Seeds were stratified for 3 days in the dark at 4 °C and transferred for 7 days to 22°C in the light (16h light /8h dark).

Plantlets of *B. distachyon* and *L japonicus* were transferred to 14 cm ConeTainers™ (2 plantlets per ConeTainer™, Stuewe & Sons Inc., Tangent Oregon, USA) containing a sand-vermiculite mix (3:1) and grown at 20°C temperature, 50% air humidity and 16-h-light/8-h-dark cycles. Each ConeTainer™ was inoculated with 140 spores of one of the four *R irregularis* strains (A4, DAOM-664343; A5, DAOM-664344; G1, DAOM-970895; SL1, DAOM-240409) that were produced in *Daucus carota* root organ culture (ROC) and extracted from the phytagel medium with 10 mM citrate buffer pH 6; and some colonized carrot roots pieces. Each cone was fertilized once a week with 10 ml of half Hoagland medium containing 20 μM phosphate and watered twice a week with distilled water *R irregularis* colonized *L. japonicus* and *B. distachyon* roots were stained with acid ink ^45^. Root length colonization was quantified using a modified gridline intersect method and 10X magnification at a light microscope. Root length colonization of *L. japonicus* was on average 89% by A5, 98% by A4, 92% by G1 and 68% by SL1; *B. distachyon* root length colonization was 64% by A5, 78% by A4, 60% by G1 and 60% by SL1 **(Supplemental Tables 5, 6)**.

### RNA extraction and sequencing from ROCs and whole plants

RNA was extracted from mycelium and colonized roots of *Rhizophagus irregularis* strains A4, DAOM-664343; A5, DAOM-664344; G1, DAOM-970895, SL1 DAOM-240409) after cultivation in symbiosis with either *Daucus carota* or *Cichorium intybus* in root organ cultures (ROCs). These contaminant-free root cultures allow for the propagation of AMF dikaryons under remarkably stable environmental conditions and thus represent an optimal system to measure epigenetic effects that are not confounded by multiple parameters (as it would be expected of in planta and pot cultures). Root colonization was measured using established protocols^45^, with 10 random root samples from the same replicate being investigated (**Supplemental Figure 11**). If the colonization was weak, or less than 6-7 root samples were strongly colonized, those plates were not used for RNA extraction.

Overall, SL1 ROC colonization was poor for *Cichorium intybus* and the RIN of RNA extracted from extraradical mycelium of this strain did not exceed our minimal sequencing threshold of 7. As such, these conditions were thus not further analyzed for this strain. In contrast, strong root colonization (80%) was visualized for all other strains and ROC plates regardless of host (**Supplemental Figure 11**). No intra-plate differences in colonization were observed, and any ROC plate with weak colonization was discarded for downstream RNA procedures. Total RNA was extracted from flash frozen colonized roots or extraradical tissue using the RNeasy plant mini kit (Qiagen) following the manufacturer recommendations and treated with DNAse. RNA quantification with the Q-bit spectrometer, and RNA quality was determined by using electrophoresis on an Agilent Bioanalyzer. Overall, samples showed an average RIN or 8.1. Four RNA biological replicates per condition (material from distinct four ROC plates) were sent for single strand, 100bp paired-end sequencing at Discovery Life Science (Huntsville, AL).

For RNAseq from *R. irregularis* colonizing *L. japonicus* and *B. distachyon*, the roots of the two plants of each ConeTainer™ were cut and divided into two equal parts for the quantification of root length colonization and for RNA extraction. For RNA extraction the root tissue was flash frozen in liquid nitrogen and kept at -80°C until use. The material was ground with mortar and pestle and the RNA was extracted with the Spectrum Plant Total RNA kit (Sigma) and treated with DNAse I amplification grade (Sigma). The efficiency of the DNAse treatment was tested by PCR. RNA concentration was measured with a Qubit 2.0 using Qubit RNA Broad Range Assay (Invitrogen). Quality and concentration were examined using Bioanalyzer RNA Nano Chips (Agilent), all samples had a RIN ≥ 7. The RNA sequencing was performed on a NovaSeq6000 (Illumina) at IMGM Laboratories GmbH (Martinsried, Germany) using NovaSeq 6000 S1 200 chemistry (Illumina) with paired end reads (2 × 101 bp) and index reads i7 and i5 (10 bp each).

RNA sequencing data were cleaned with fastp 0.22.0 and default settings ^46^. Salmon v.1.8.0 ^47^ was used to align the clean RNA-seq reads to the transcripts and to estimate transcript abundances in each sample (salmon index –keepDuplicates and salmon quant –validateMappings). The entire genome was used as a decoy sequence during the read mappings. We used Tximport and DESeq2 ^48^ to assess gene differential expression (*P*adj < 0.1).

### Chromosome assembly and annotation

The HiFi reads of the strains A4, A5 and G1 were assembled with hifiasm 0.16.1 in Hi-C integration mode and with default parameters ^49^. For G1 --hom-cov was set to 181. The resulting contigs were scanned for possible contaminations and mitochondrial sequences by running a Diamond blastx (diamond v0.9.14.115, Database format version = 1) and flagged contigs were removed from the assembly ^50^. Phasing of the assembled haplotypes was confirmed with the NuclearPhaser pipeline (MAPQ=10, 100 Kb Hi-C contact map resolution) ^22^. For scaffolding, the Hi-C reads were first mapped to each haplotype using BWA-MEM 0.7.17 ^51^ and further processed according to the Arima Genomics pipeline (https://github.com/ArimaGenomics/mapping_pipeline/blob/master/01_mapping_arima.sh).

Scaffolding was performed using SALSA 2.2 ^52^. Hi-C contact maps were produced with HiC-Pro 2.11.1 ^53^ and visually examined for correct scaffolding.

De novo repeats were predicted with RepeatModeler 2.0.0 and the option -LTRStruct ^54^. These were merged with the RepeatMasker repeat library and RepeatMasker 4.1.0 was run with this combined repeat database (http://www.repeatmasker.org).

Gene annotation was achieved by using Funannotate (v.1.7.4)

(https://zenodo.org/record/2604804#.YPm336iSnIU) on repeat-masked genome assemblies. Genome annotations were performed using RNA-seq reads and protein models available for strains A4 and A5 (JGI, ^29^), and using RNA-seq for strain G1 and RNA-seq reads and protein models available for strains A4 were used for SL1. The quality of annotations was evaluated using the Benchmarking Universal Single-Copy Orthologs software v5.2.2 ^55^. Conserved protein domains were predicted using Pfam v.27 ^56^. SignalP 4.1 (-t euk -u 0.34 -U 0.34) ^57^ and Tmhmm 2.0 ^58^ were used to predict secreted proteins. A protein was called secreted if it was predicted to have a signal peptide but no transmembrane domains. Effector candidates were predicted with EffectorP 3.0 ^59^.

To assess differences in repeat content, transposable element locations were extracted from the RepeatMasker output file and simple repeats, unknown and low-complexity repeats, satellites, tRNA, sRNA and rRNAs were fremoved. PfamScan ^56^ was used with default parameters to identify the protein domain annotations in all strains. To create the Pfam heatmaps, the domain numbers were counted for each *R. irregularis* strain or A/B compartment, and *t-*tests were used to highlight the protein domain abundances for each category.

We used dnadiff from the MUMmer suite for structural variation calculations between strains and compartments^60^. D-Genies dot plots were also used to analyze overall synteny between haplotypes and pinpoint structural variation sites between them ^61^. Orthology analyses were made by OrthoMCL ^62^ using the following parameters: 50% identity and 50% coverage using protein sequences of all five assemblies. All karyoplots were produced using KaryoploteR ^63^.

GO annotations were performed using EggNOG (v5.0.2), with the funog database ^64^. Proteins from all genes and 1-to-1 orthologous groups upregulated across conditions were selected and their GO annotations were analyzed. GO classifiers were categorised using WEGO 2.0 ^65^, and the results visualized using R.

### Identification of chromosome compartments and topologically associated domains

We called A/B compartments with Hicexplorer 3.6^66^ using the command hicPCA. Inter- and intrachromosomal Hi-C contact maps were produced with HiC-Pro 2.11.1 (MAPQ = 10) and Hicexplorer 3.6 and these Hi-C contact maps were manually inspected to assign chromosomal regions to A/B compartments. Specifically, the regions along each chromosomes carrying the same PCA1 values – that is, positive or negative – were assigned to the same compartment. Following this assignment, compartments were manually inspected to investigate their proximity with other chromosomes. Those in close physical proximity with the smallest chromosomes were assigned to compartment A, while those carrying the counterpart PCA1 signal were assigned to compartment B. This process was manually repeated for all 32 chromosomes individually to separate the chromosomal regions into separate A/B compartments. Bedtools was used to assess overlap between genomic features such as genes/repeats and compartments ^67^. The strength of the PCA1 values for the strains G1 and SL1 allowed the identification of compartment for only a subset of the chromosomes.

### Phylogenetic analyses

The haplotype genomes from AMF heterokaryons and ddRAD-seq data were used included to perform a phylogenetic of R. irregularis populations using the Phame pipeline ^68^. The ddRAD-seq data from 61 *R. irregularis* isolates ^28^ (all available replicates included) were quality assessed using FASTQC (https://www.bioinformatics.babraham.ac.uk/projects/fastqc/), and Illumina adapters were trimmed using TagCleaner ^69^. Reads containing uncalled bases (N) and shorter reads (<50bp) were removed. The demultiplexing of the dataset was achieved with the *process_radtags* from the Stacks pipeline v2.60 ^70^.

To construct the phylogenetic tree, the 8 haplotype genome assemblies as well as five chromosome-level homokaryotic assemblies ^25^ and the ddRAD-seq reads that were processed with PhaME (version 1.0.2) using these parameters: data = 4, reads = 2, code = 0, cutoff = 0.0001, reference = 1, reffile = DAOM-197198 genome assembly ^68^. IQTREE algorithm was used with GTR+FO mode and1000 bootstrap replicates on the 105,979 bp alignment file to construct the phylogenetic tree^62^. The phylogenetic tree was then visualized with FigTree (http://tree.bio.ed.ac.uk/software/figtree/). Admixture (v1.3.0) was used for cross-validation of the *K* values and the *K* value with the lowest coefficient of variant was selected (*K* = 11) ^71^. Nucleotide alignment file created by Phame was then used by NGSadmix with -K 11 parameter was used to create the admixture plots ^72^.

### PCR amplification and digital droplet PCR (ddPCR)

PCR amplification and digital droplet PCR (ddPCR) were done on each sample in order to identify the nuclear ratio of the MAT locis for each strain. The PCR mixture for each sample contained 12 µL of 2X Supermix for Probes (No dUTP) (Bio-Rad), 1.2 µL of each primers/probe mixture (PrimeTime® Std qPCR Assay (500 rxn), 7.2 µL of autoclaved dH2O and 1 µL of approximately 50 ng/µL of extracted DNA. The following replicates were used for ddPCR: A5/Chicory, 23 replicates; A5/P68, 24 replicates; A4/Chicory, 24 replicates; A4/P68, 24 replicates, G1/Chicory, 22 replicates, G1/P68, 18 replicates.

The MAT-1 (FAM) primers 5’-CATCAACAAGTCAACGATTTAT-3’ and 5’-GTGGATACATGACATGGTGT-3’ and probe 5’-CAGAAACATTTAATAATAATAATACACGTT-3’; MAT-2 (HEX) primers 5’-TACACAACAAGTCAACGATG-3’ and 5’-CATGATGCTCAATATTAAGTG-3’ and probe 5’-GTAATGAAATTATAGAAGGAAATATTAG-3’; MAT-3 (HEX) primers 5’-CGTCAAAGAATCACGACACTC-3’ and 5’-CATTATTCACAATTGCGTTCGG-3’ and probe 5’-CATATAAGAAACAAAAATGCCTTAATTCAAG-3’; MAT-5 (HEX) primers 5’-CAGATTTAGACAAAGATATTCG-3’ and 5’-TCCTTATTACATATTTCTACA-3’ and probe 5’-CTTGTATATCAACTGTAACATATCAG-3’; MAT-6 (FAM) primers 5’-CCGTCAAAGACTCACAACAC-3’ and 5’-CATTATTCACAATTGCGTTCGG-3’ and probe 5’-CACATCCGTATATATGTGCAGATAAACAA-3’ were used for A4 (MAT-1 and MAT-2), A5 (MAT-3 and MAT-6) and G1(Mat-1 and MAT-5) accordingly ^1^. Five to twelve PCR replicates were used depending on availability for each condition. Two positive controls (mycelium extracted DNA) and two no-template controls (containing the two respective MAT primers/probe) were used per strain. Samples were emulsified using the Bio-Rad QX200™ Droplet Generator with 72µL of droplet Generation Oil for Probes per sample. Samples were transferred to a 96-well plate for PCR and the C1000 Touch™ Thermal Cycler (Bio-Rad) was used at the following cycling conditions: 10 minutes at 95°C, 1 minute at 94°C, 2 minutes at 52.5°C, followed by 44 cycles of 1 minute at 94°C and 2 minutes at 52.5°C, then 10 minutes at 98°C and an infinite hold at 4°C. At every step, a ramp rate of 1°C/second was used ^1^. Following PCR amplification, ddPCR and absolute DNA quantification analysis were performed using the Bio-Rad QX200™ Droplet Reader. The droplet reader identifies each droplet as positive or negative in regard to the fluorescence level of the dye probe, in this case FAM or HEX. Dual-target detection for each sample via multiplexing was used (detection of the 2 respective MAT locis within the same well) to measure the ratio of the MAT-loci within each sample. The software QuantaSoft Analysis Pro (1.0.596) fits the fraction of positive droplets into a Poisson algorithm to determine the initial absolute concentration of DNA. The positive and no-template controls were used as guides to manually set the threshold using the Threshold Line Mode Tool at the 2D amplitude mode of QuantaSoft Analysis Pro (1.0.596). Wells containing less than 10,000 droplets were excluded from the analysis.

### SNP calling and plotting allelic frequencies of haplotypes

PacBio reads produced from each heterokaryotic strain in this study were mapped individually to haplotype genome assemblies using minimap2 (v2.22-r1101) using –MD –ax map-hifi parameters ^73^. Variation calling was performed with Freebayes (v1.3.6) using freebayes-parallel, with default parameters ^74^. Bcftools filter was used to filter, SNPs based on read depth (maximum read depth: DP < 1.25× genome mean coverage; minimum read depth: DP > 0.75× genome mean coverage and quality QUAL> 20 ^75^. Only SNPs that are found on 1-to-1 alignment regions produced by dnadiff between haplotypes were analysed, and SNPs on repeat regions were removed. These final bedfiles were processed in an R script to plot SNP frequency (AO/DP ratio) histograms and density plots for each SNP positions.

## Acknowledgments

We thank Pierre-Marc Delaux, Timothy James, Allyson MacLean and Vasilis Kokkoris for their helpful comments on an earlier version of this manuscript. Our research is funded by the Discovery program of the Natural Sciences and Engineering Research Council (RGPIN2020-05643), a Discovery Accelerator Supplements Program (RGPAS-2020-00033). N.C. is a University of Ottawa Research Chair in Microbial Genomics. J.S. was supported by an Australian Research Council (ARC) Discovery Early Career Researcher Award (DE190100066).

## Data accessibility

All genome and RNA-seq data and reads newly obtained are available in GenBank under the BioProject PRJNA922099.

