## Supplemental Figures for "Resolving the haplotypes of arbuscular mycorrhizal fungi highlights the role of two nuclear populations in host interactions"

A)

B)

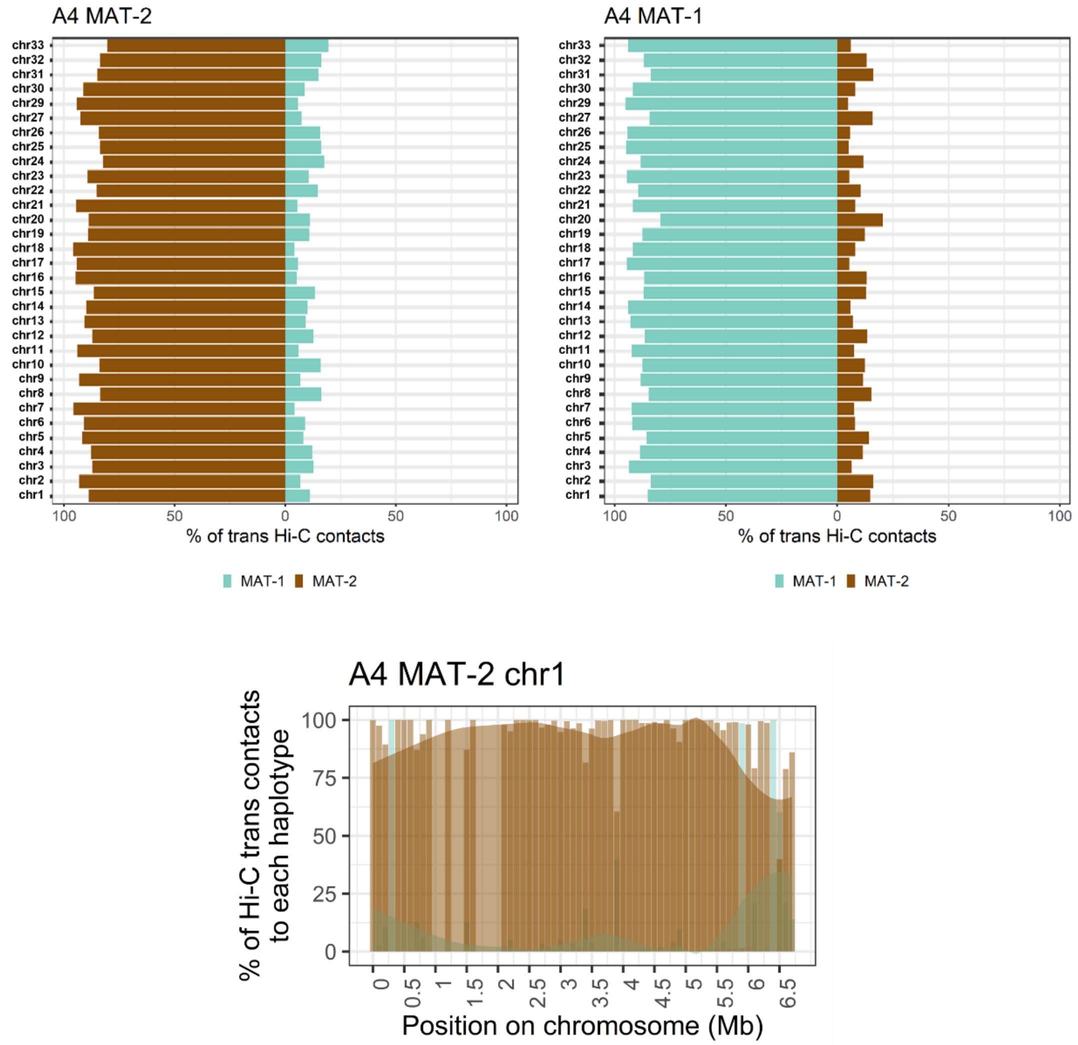


**Supplemental Figure 1: The Hi-C contacts between nuclear-separated haplotypes of heterokaryotic *Rhizophagus irregularis* A4 strain. A) Percentage of Hi-C contacts that link within and between nuclear-separated haplotypes.** The assemblies have a strong dikaryotic Hi-C phasing signal, with over 90% of inter-chromosomal (trans) contacts occurring within the nucleus. **B) Graphs showing the % of Hi-C trans contacts that link to the two haplotypes for chromosome 1 of MAT-2 haplotype.** This chromosome is fully-phased, with only weak Hi-C trans contact to the other nucleus in some regions.

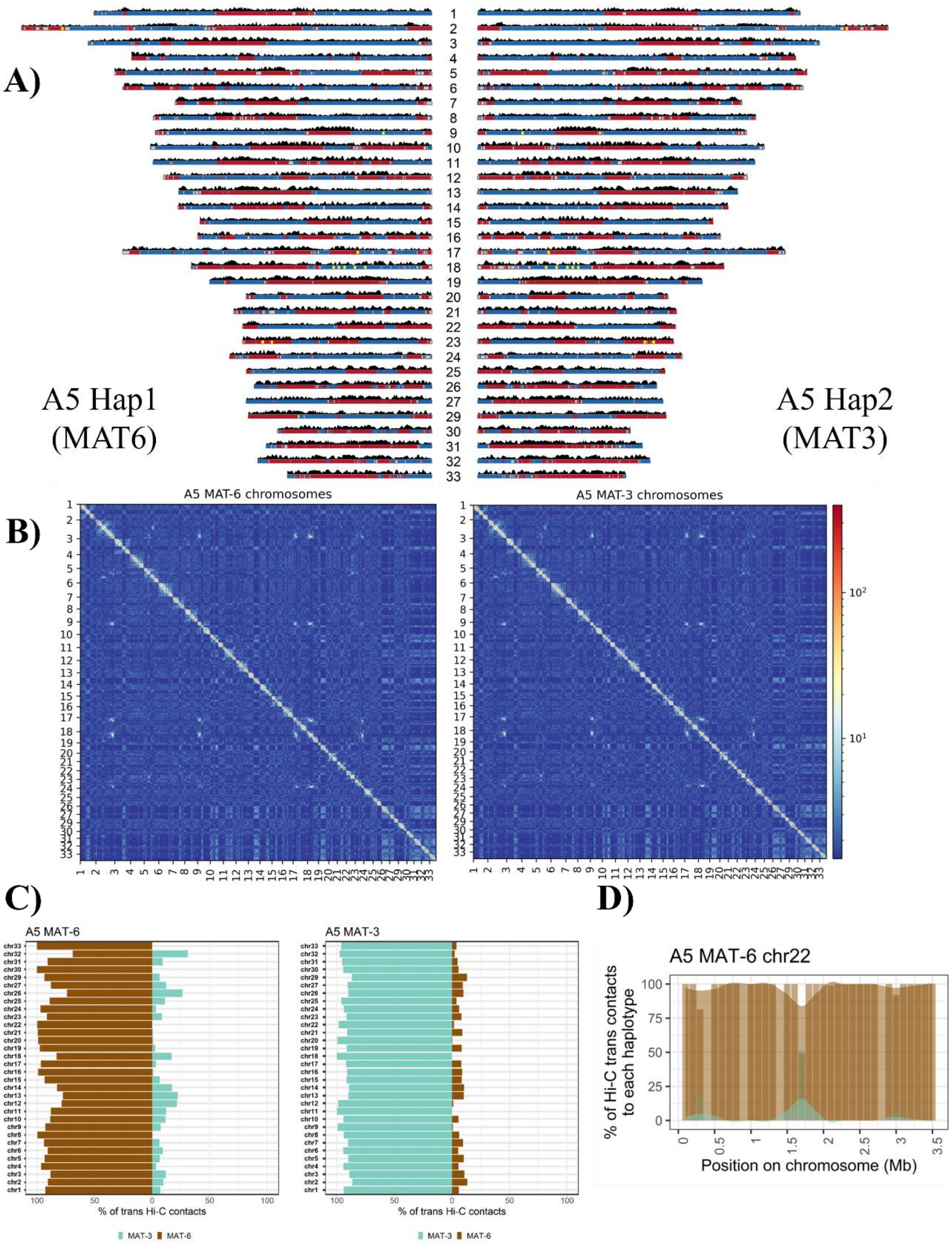


**Supplemental Figure 2: The phased chromosomes of the AMF heterokaryotic strain A5. A) The karyoplots represent the chromosomes of the haplotypes.** Gene density is shown in black (20 Kb windows). A and B compartments are shown in red and blue, respectively. **(B) The Hi-C contact map of the haplotypes show a compartmentalization of the chromosomes into A and B compartments. (C) Percentage of Hi-C contacts that link within and between nuclear-separated haplotypes.** The assemblies have a strong dikaryotic Hi-C phasing signal, with over 90% of inter-chromosomal (trans) contacts occurring within the nucleus. **(D) Graph showing the % of Hi-C trans contacts that link to the two haplotypes for chromosome 22 of MAT-6 haplotype.** The chromosomes are fully-phased, with only weak Hi-C trans contact to the other nucleus in some regions.

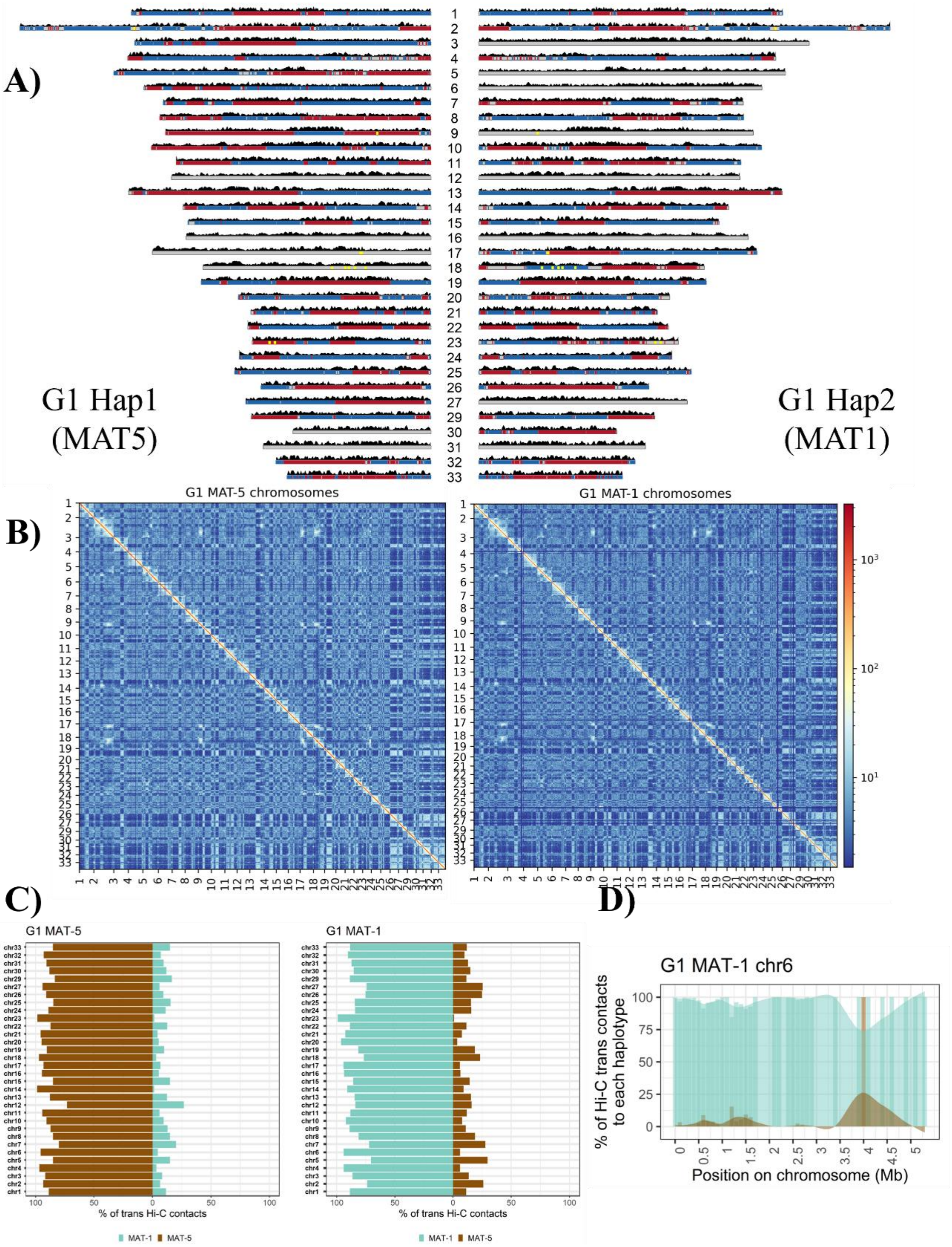


**Supplemental Figure 3: The phased chromosomes of the AMF heterokaryotic strain G1. A) The karyoplots represent the chromosomes of the haplotypes.** Gene density is shown in black (20 Kb windows). A and B compartments are shown in red and blue, respectively. **(B) The Hi-C contact map of the haplotypes show a compartmentalization of the chromosomes into A and B compartments. (C) Percentage of Hi-C contacts that link within and between nuclear-separated haplotypes.** The assemblies have a strong dikaryotic Hi-C phasing signal, with over 90% of inter-chromosomal (trans) contacts occurring within the nucleus. **(D) Graph showing the % of Hi-C trans contacts that link to the two haplotypes for chromosome 6 of MAT-1 haplotype.** The chromosomes are fully-phased, with only weak Hi-C trans contact to the other nucleus in some regions.

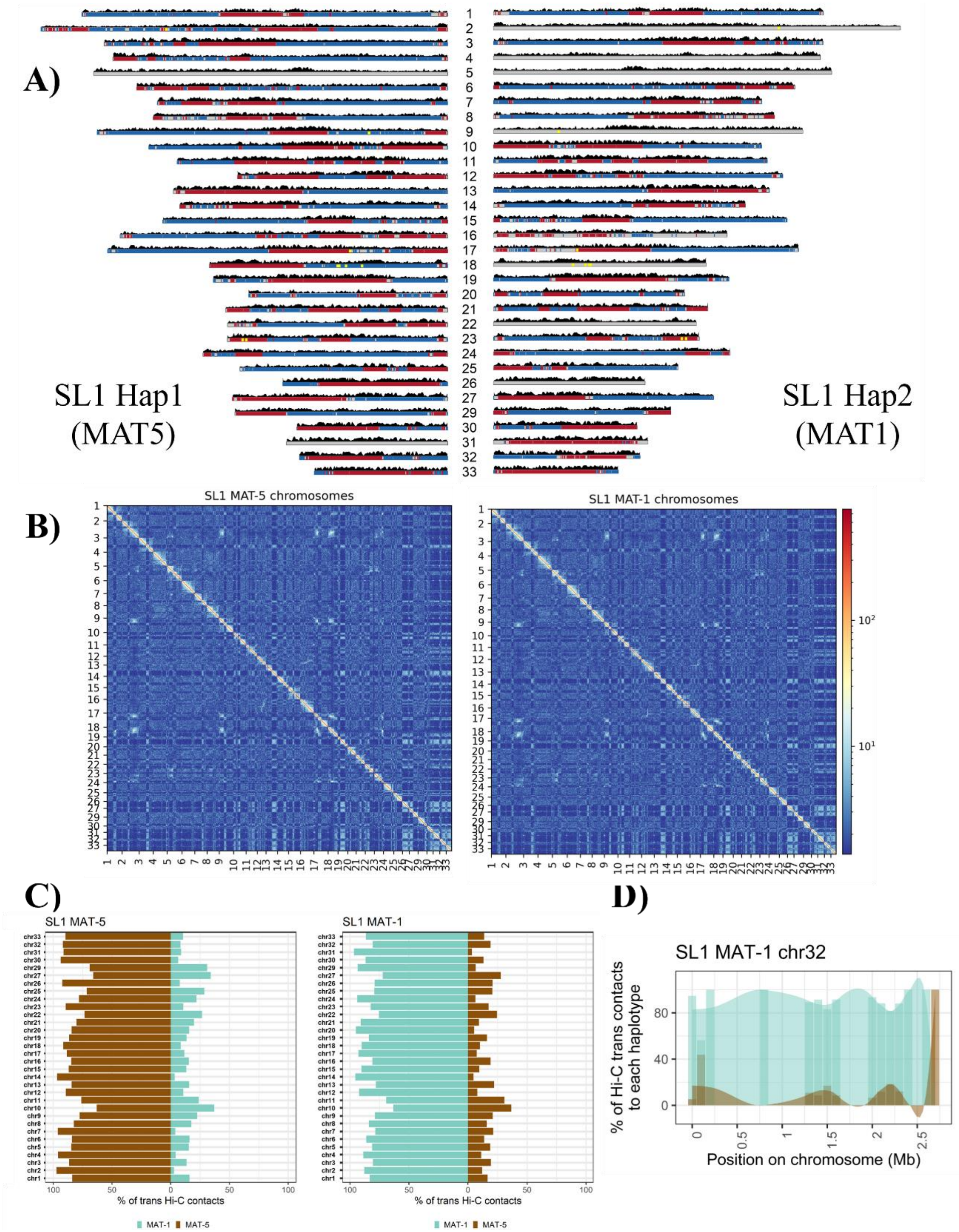


**Supplemental Figure 4: The phased chromosomes of the AMF heterokaryotic strain SL1. A) The karyoplots represent the chromosomes of the haplotypes.** Gene density is shown in black (20 Kb windows). A and B compartments are shown in red and blue, respectively. **(B) The Hi-C contact map of the haplotypes show a compartmentalization of the chromosomes into A and B compartments. (C) Percentage of Hi-C contacts that link within and between nuclear-separated haplotypes.** The assemblies have a strong dikaryotic Hi-C phasing signal, with over 90% of inter-chromosomal (trans) contacts occurring within the nucleus. **(D) Graph showing the % of Hi-C trans contacts that link to the two haplotypes for chromosome 32 of MAT-1 haplotype.** The chromosomes are fully-phased, with only weak Hi-C trans contact to the other nucleus in some regions.

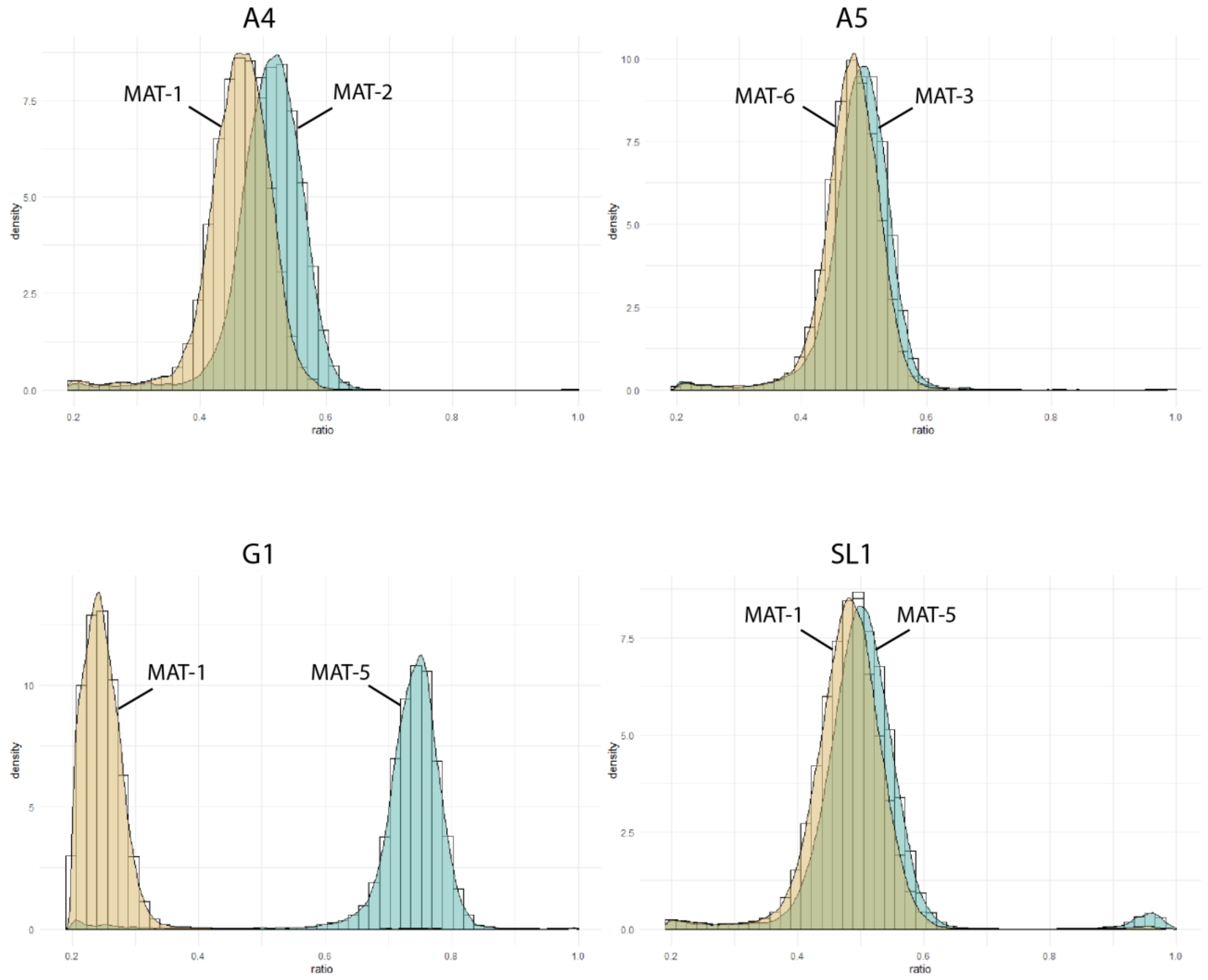


**Supplemental Figure 5: Histograms of allele frequency distribution of *R. irregularis* strains A4, A5, G1 and SL1 haplotypes.** The histograms of allele frequency distribution are based on bi-allelic SNPs filtered based on the relative genome coverage of each haplotype to exclude contaminants and are overlapped by density curves (black). The X and Y axes represent the SNP frequency (ratio) and density, respectively. Analyses are based SNPs identified along 1-1 aligned regions (excluding repetitive regions), representing between 64,331,134 bp to 69,274,838 bp of each haplotype in total. These SNP reflect relative abundance in extraradical of Carrot *cv.* P68.

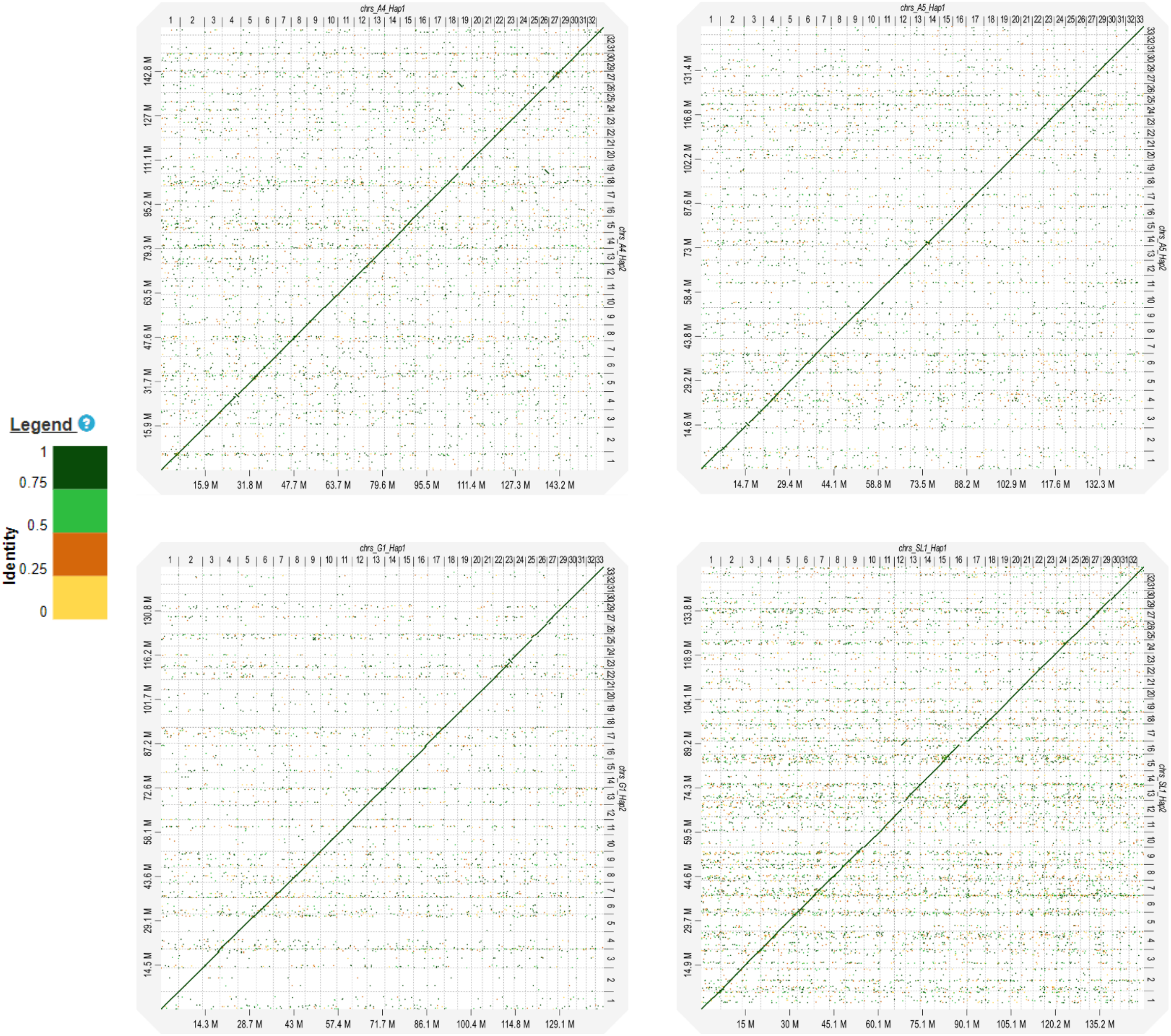


**Supplemental Figure 6: D-Genies dot plots show overall synteny and reveal few structural variations between coexisting haplotypes in heterokaryotic strains.** Haplotypes from the same strains are located in x- and y-axes. The aligned regions are represented as dots, and the colors represent identity. The diagonal lines represent synteny between two haplotypes. Breaks indicate rearrangement events.

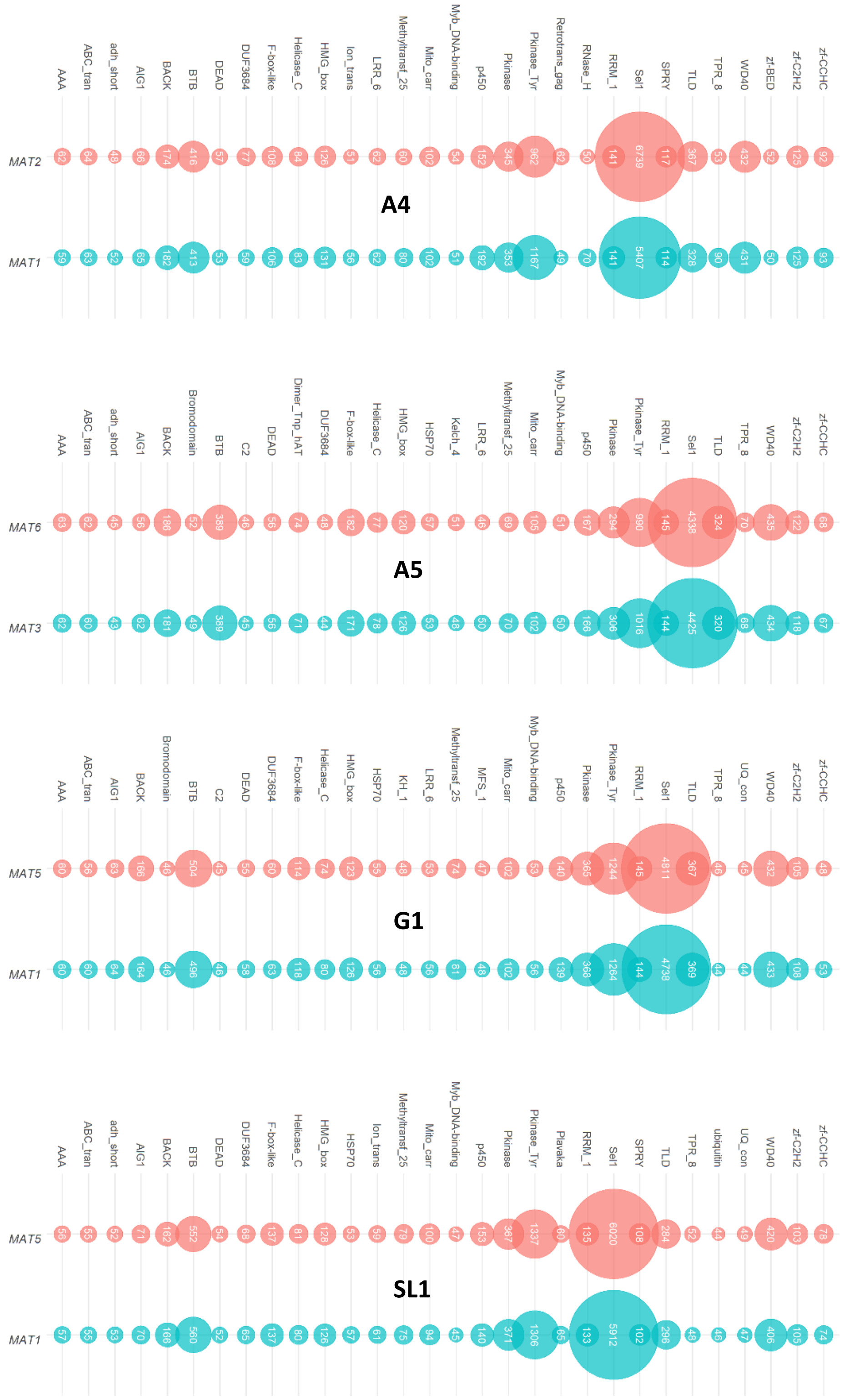


**Supplemental Figure 7: Haplotypes differ from each other in protein content. Pfam domain number comparisons of the heterokaryotic *Rhizophagus irregularis* strain haplotypes.** The Pfam domain numbers are shown in bubbleplots for each strain. 30 most abundant pfam domains are shown, the strain names are placed in the center of the bubbleplots. The MAT loci of different haplotypes are shown in different colors.

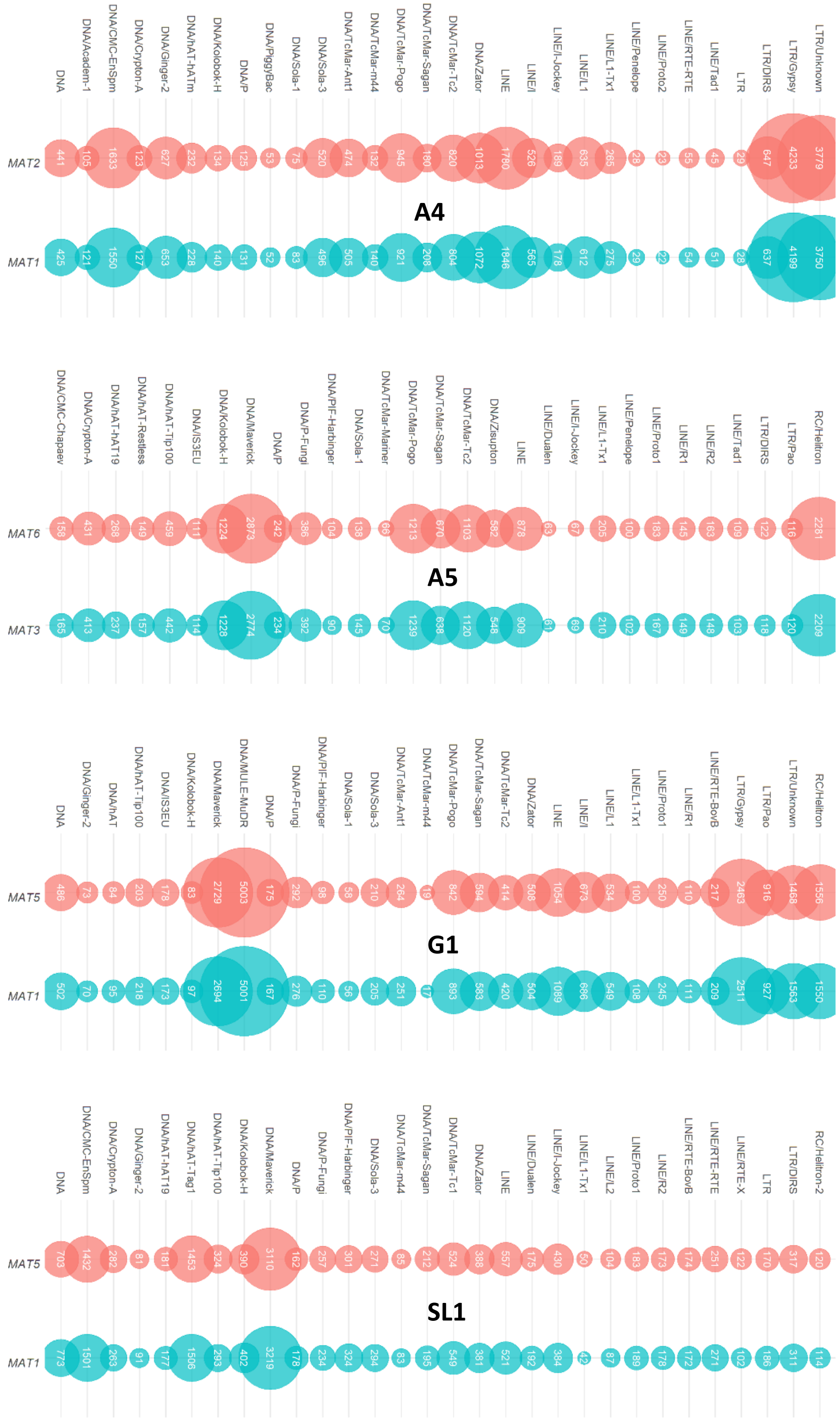


**Supplemental Figure 8: Haplotypes differ from each other in repeat content. Transposable element number comparisons of the heterokaryotic *Rhizophagus irregularis* strain haplotypes.** The repeat numbers are shown in bubbleplots for each strain. 30 transposable elements with the most diverging numbers are shown, the strain names are placed in the centre of the bubbleplot. The MAT loci of different haplotypes are shown in different colors.

**A)**

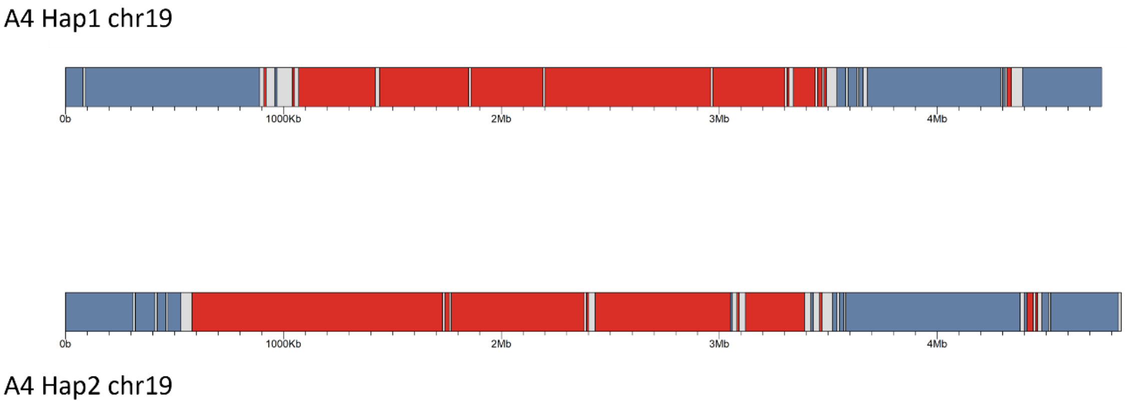


**B)**

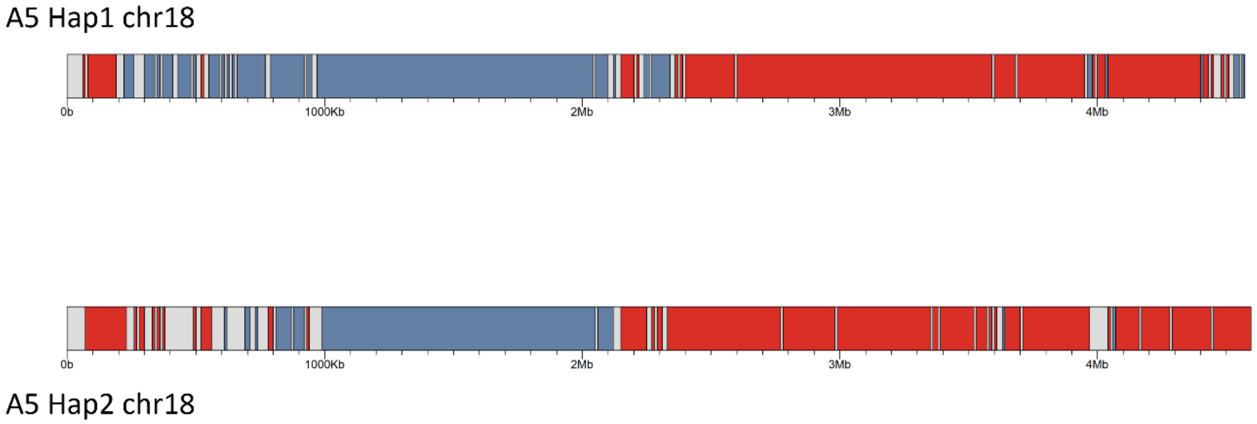


**Supplemental Figure 9: Karyoplots representing compartment variability between two homologous chromosomes of *Rhizophagus irregularis* heterokaryotic strains.** The length of the karyoplots represent the chromosome sizes, red color represents compartment A and blue color represents compartment B. Grey areas do not belong to a compartment. Pink ribbons represents linear synteny between chromosomes, yellow ribbons indicate inverted synteny. **A) Karyoplots representing compartment variability in chromosome 19 of *Rhizophagus irregularis* A4 strain. B) Karyoplots representing compartment variability in chromosome 18 of *Rhizophagus irregularis* A5 strain.**

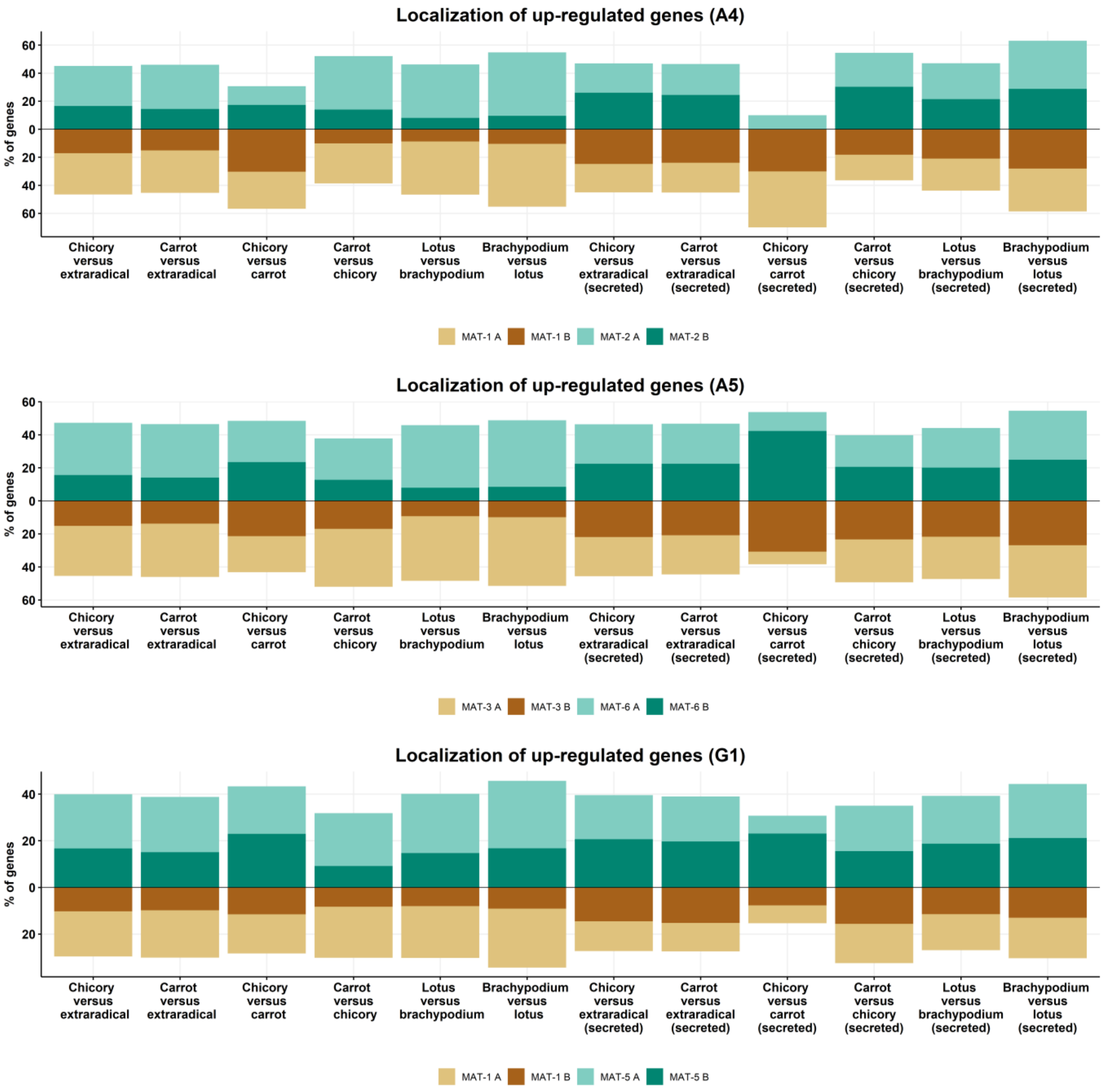


**Supplemental Figure 10: Chromosome localization of gene upregulation**. When all differentially regulated genes are considered, upregulation of secreted genes is higher in planta in the B-compartment, and in some cases preferentially affects one haplotype.

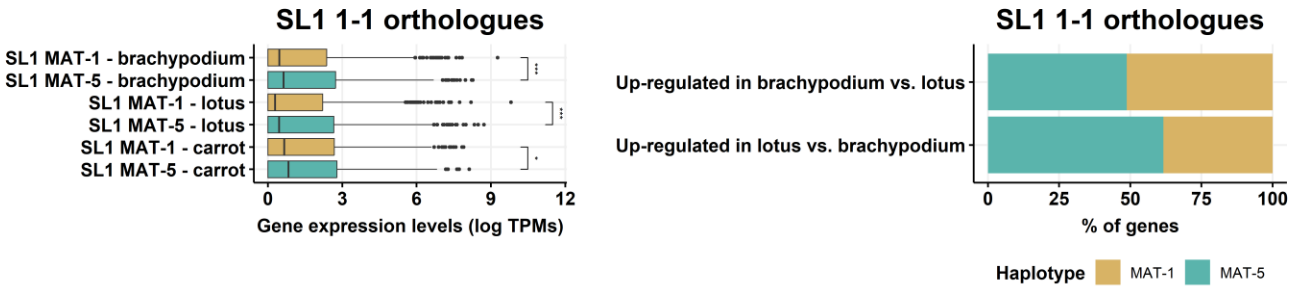


**Supplemental Figure 11: Haplotype specific RNA-seq analyses SL1.** A. Haplotype-specific expression (left) and regulation of 1-1 orthologues (right). Left: Box plot edges show the first and third quartile in box plots, with the median shown as the middle lane. The data range is indicated by whiskers and dots represent outliers. Asterisks indicate a t-test statistical significance at least P < 0.05. Right: % of genes upregulated in each haplotype across the conditions used in this study.

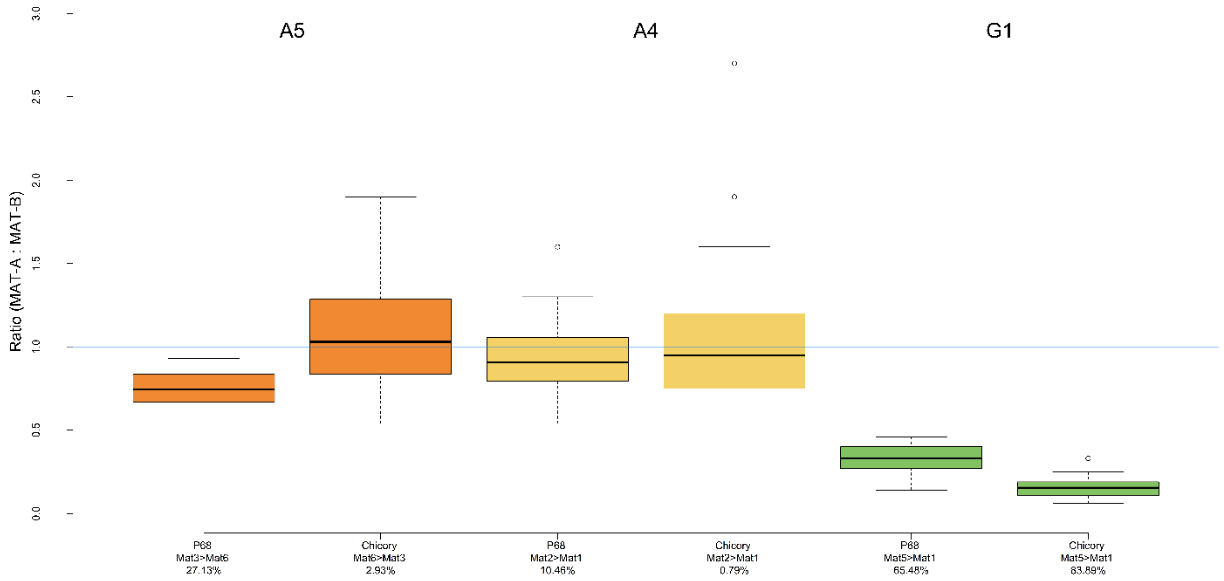


**Supplemental Figure 12: Change in relative nucleotype abundance due to host in dikaryon AMF strains A4, A5 and G1.** The ratio between both nucleotypes A and B (FAM versus HEX) due to either the host *Daucus carota* or *Cichorium intybus* of strains A4, A5 and G1 was measured. The blue line indicates a 1:1 ratio. Dominance of nucleotype A is present when the boxplot is above the blue line, and dominance of nucleotype B is the case when the boxplot is under the blue line. The third quartile (edge of box), first quartile (edge of box), median (middle blue line), outliers (dots) and range of data (whiskers) are shown. The dominant MAT loci and percentage difference between nucleotypes per strain was calculated with the following formula: ([A/B]/A) X 100, A and B is the mean number of the most and least abundant nucleotype respectively. The following replicates were used for ddPCR: A5/Chicory, 23 replicates; A5/P68, 24 replicates; A4/Chicory, 24 replicates; A4/P68, 24 replicates, G1/Chicory, 22 replicates, G1/P68, 18 replicates.

A4

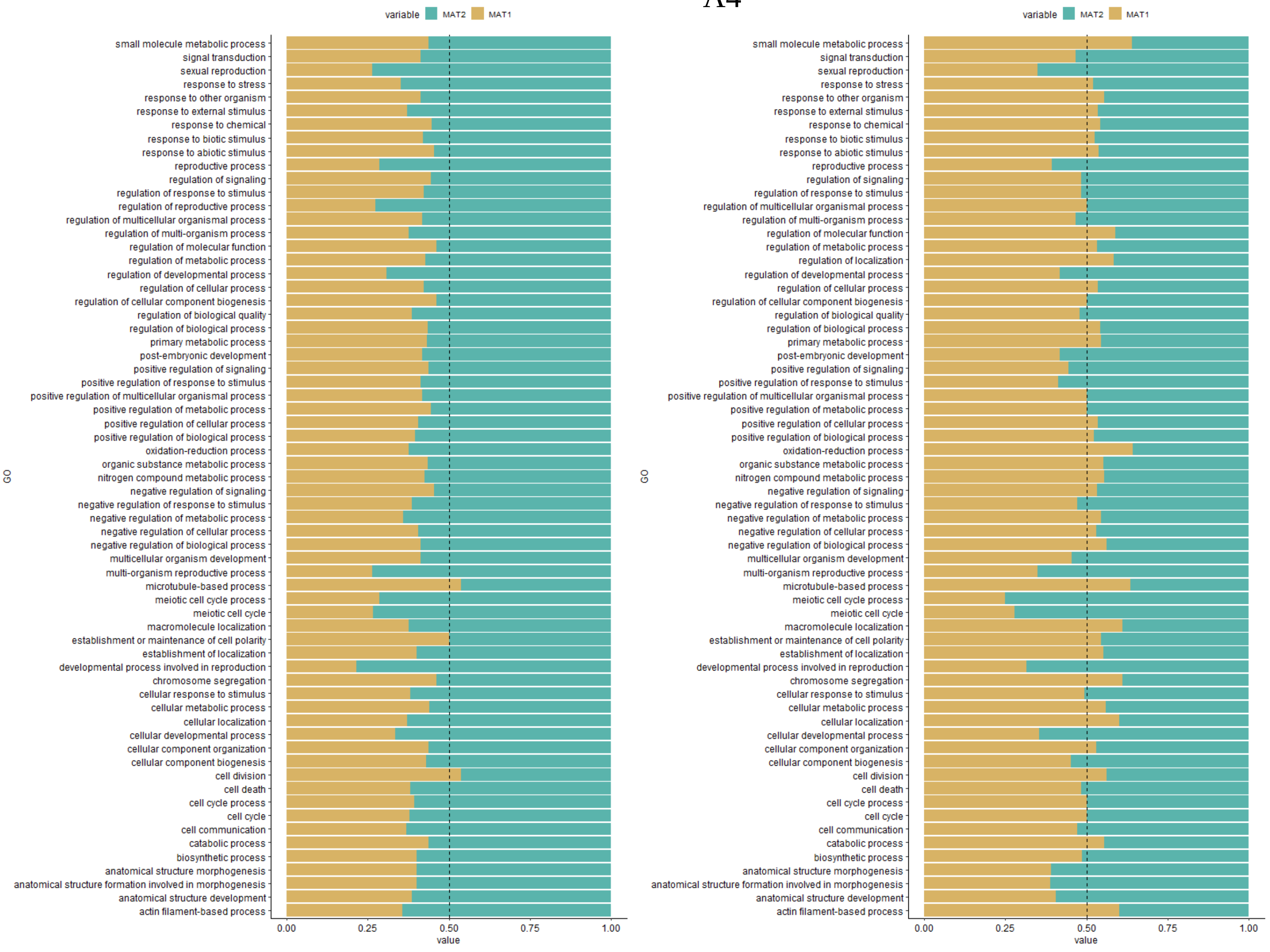


In planta *C. intybus* vs extraradical mycelium

In planta *D. carota* vs extraradical mycelium

A5

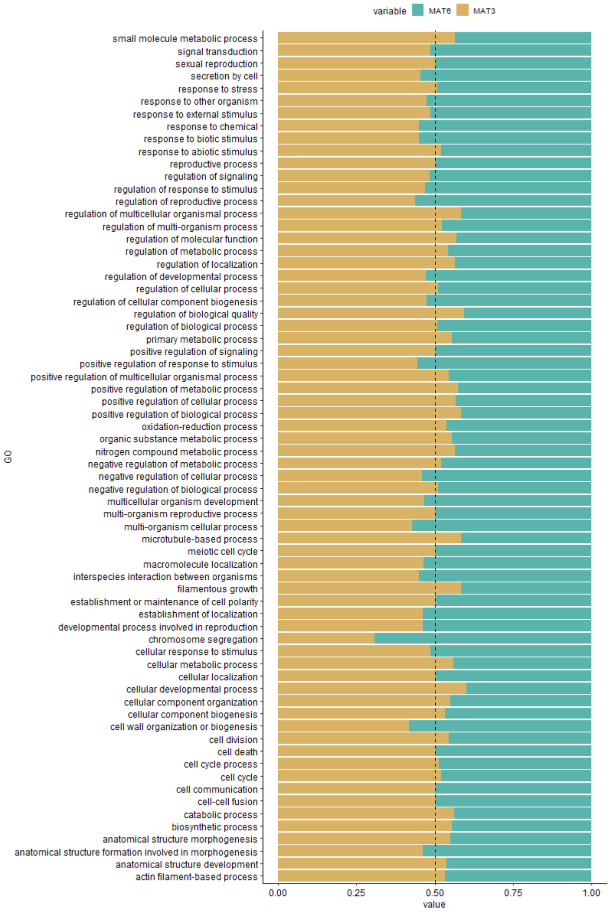


In planta *C. intybus* vs extraradical mycelium

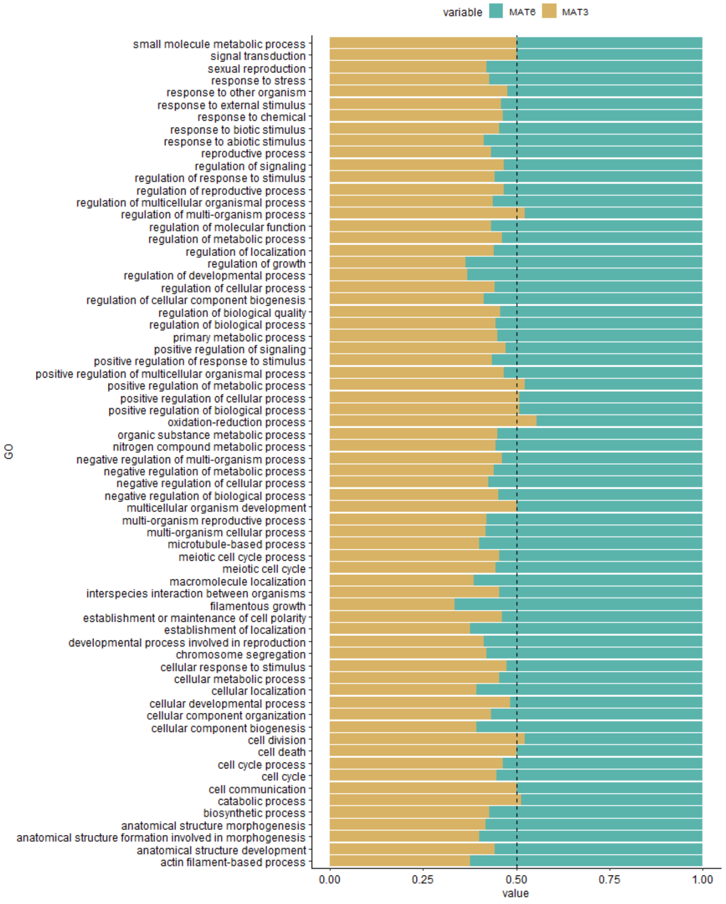


In planta *D. carota* vs extraradical mycelium

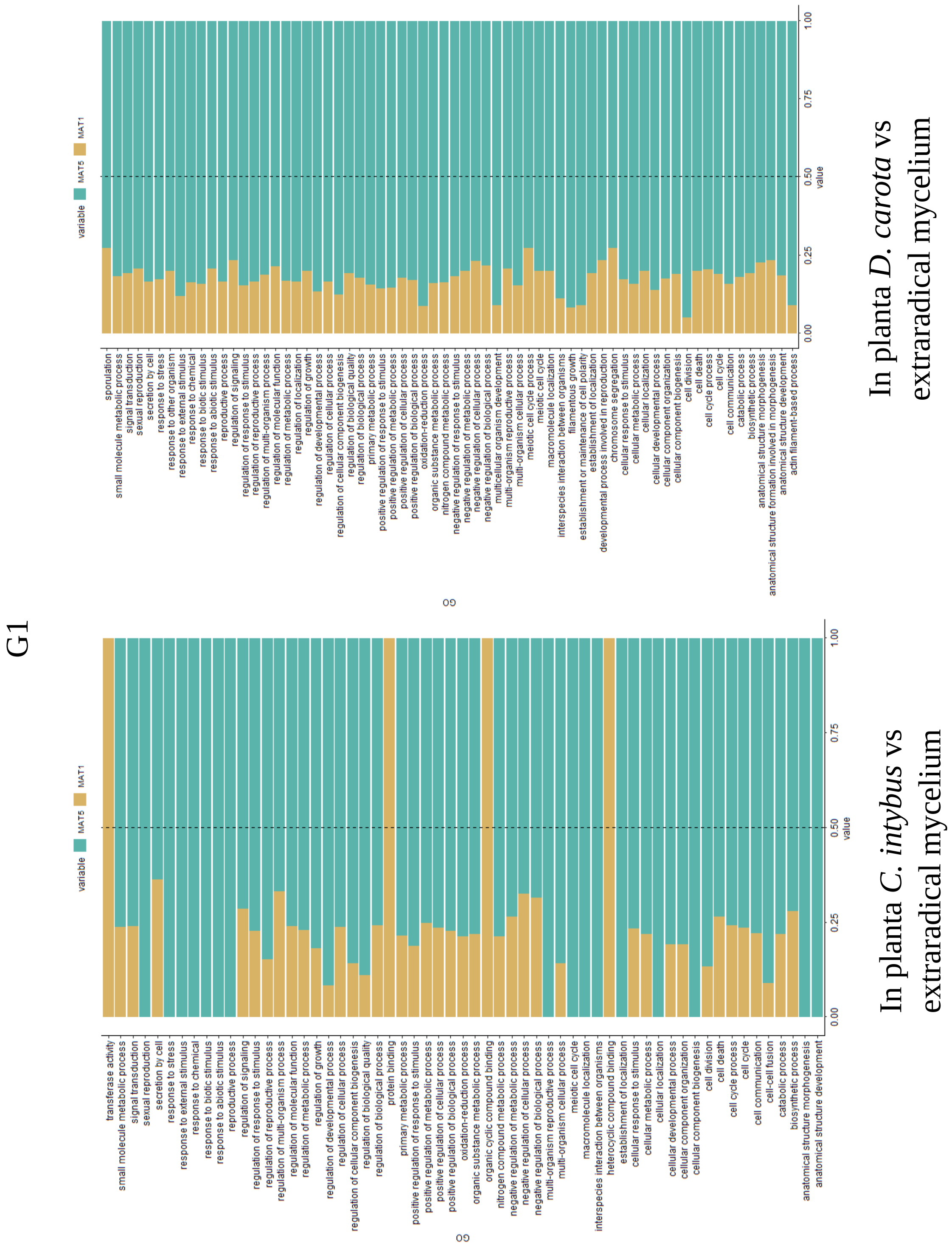


**Supplemental Figure 13: The ratio of upregulated genes and their GO annotations.** Upregulated genes for each GO terms were analyzed to reveal the contribution of each haplotype. In planta *C. intybus* and *D. carota* expressions were compared to expression of extraradical spores and mycelium. Haplotypes are shown in different colors. 0.5 ratio is shown with dashed line.

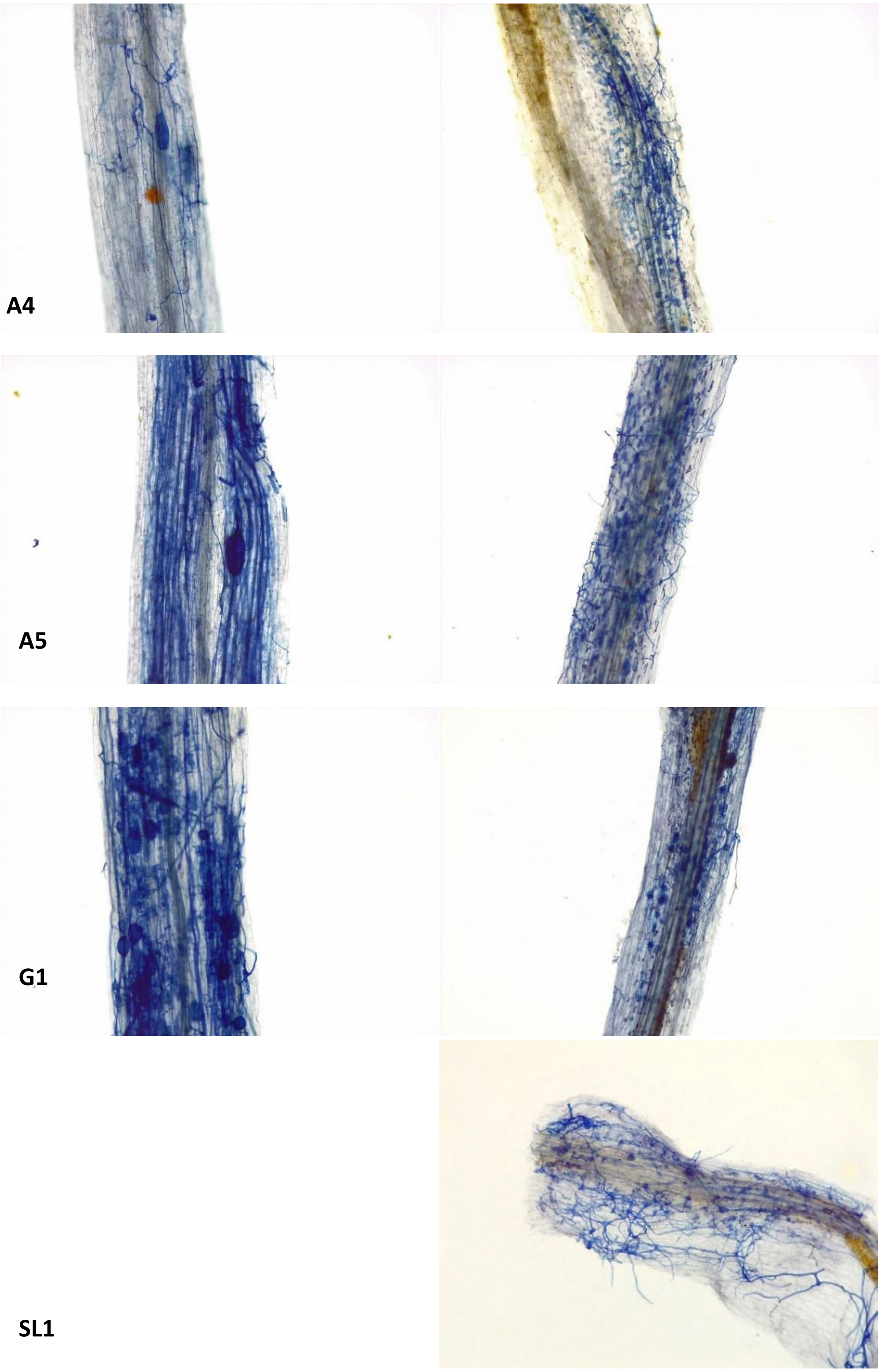


**Supplemental Figure 14: Microscopic observations of arbuscules and spores of *Rhizophagus irregularis* heterokaryotic strains following black Shaeffer staining of the colonized roots.** The images on the left side show colonized *Daucus carota cv* P68 roots and the images on the right side show colonized *Cichorium intybus* roots. The name of the strain colonizing these roots are indicated on the left. Images are representatives of root colonization for each strain for each host and within each replicate.
