## Supplemental Tables for "Resolving the haplotypes of arbuscular mycorrhizal fungi highlights the role of two nuclear populations in host interactions"

**Supplemental Table 1: rRNA operon locations in heterokaryotic strains.**

| No of operon copies in genome | Assigned operon Id | location | strain |
|---|---|---|---|
| 9 | operon 1 | chr23_A4_Hap1:3948972-3954773 | A4 hap1 |
| | operon 2 | chr23_A4_Hap1:4033538-4027737 | A4 hap1 |
| | operon 3 | chr17_A4_Hap1:1468375-1462560 | A4 hap1 |
| | operon 4 | chr18_A4_Hap1:1646399-1640580 | A4 hap1 |
| | operon 5 | chr18_A4_Hap1:2012308-2006481 | A4 hap1 |
| | operon 6 | chr18_A4_Hap1:1957160-1962989 | A4 hap1 |
| | operon 7 | chr18_A4_Hap1:1851467-1857283 | A4 hap1 |
| | operon8 | chr2_A4_Hap1:6786257-6792080 | A4 hap1 |
| | operon9 | chr9_A4_Hap1:1106656-1100852 | A4 hap1 |
| 9 | operon1 | chr23_A4_Hap2:3825018-3830819 | A4 hap2 |
| | operon2 | chr23_A4_Hap2:3909584-3903783 | A4 hap2 |
| | operon3 | chr17_A4_Hap2:1439838-1434023 | A4 hap2 |
| | operon4 | chr18_A4_Hap2:1660009 -1654191 | A4 hap2 |
| | operon5 | chr18_A4_Hap2:1818184-1823998 | A4 hap2 |
| | operon6 | chr18_A4_Hap2:1951508-1957335 | A4 hap2 |
| | operon7 | chr18_A4_Hap2:2017679-2011852 | A4 hap2 |
| | operon8 | chr2_A4_Hap2:6696150-6701973 | A4 hap2 |
| | operon9 | chr9_A4_Hap2:1179831-1174028 | A4 hap2 |
| 11 | operon1 | chr23_A5_Hap1:3040099-3045898 | A5 hap1 |
| | operon2 | chr23_A5_Hap1:3217915-3212116 | A5 hap1 |
| | operon3 | chr18_A5_Hap1:1764572-1770392 | A5 hap1 |
| | operon4 | chr18_A5_Hap1:1868777-1862957 | A5 hap1 |

| | | | |
|---|---|---|---|
| | operon5 | chr18_A5_Hap1:1462146-1467979 | A5 hap1 |
| | operon6 | chr18_A5_Hap1:1664019-1658186 | A5 hap1 |
| | operon7 | chr18_A5_Hap1:1271000-1265167 | A5 hap1 |
| | operon8 | chr17_A5_Hap1:1415295-1409480 | A5 hap1 |
| | operon9 | chr2_A5_Hap1:7017538-7011709 | A5 hap1 |
| | operon10 | chr9_A5_Hap1:928574-922772 | A5 hap1 |
| 10 | operon1 | chr23_A5_Hap2:3117778-3123577 | A5 hap2 |
| | operon2 | chr23_A5_Hap2:3281085-3275286 | A5 hap2 |
| | operon3 | chr18_A5_Hap2:1773370-1779190 | A5 hap2 |
| | operon4 | chr18_A5_Hap2:1877575-1871755 | A5 hap2 |
| | operon5 | chr18_A5_Hap2:1470943-1476776 | A5 hap2 |
| | operon6 | chr18_A5_Hap2:1672817-1666984 | A5 hap2 |
| | operon7 | chr18_A5_Hap2:1277461-1271628 | A5 hap2 |
| | operon8 | chr17_A5_Hap2:1326027-1320212 | A5 hap2 |
| | operon0 | chr2_A5_Hap2:6882018-6876189 | A5 hap2 |
| | operon10 | chr9_A5_Hap2:840101-834299 | A5 hap2 |
| 11 | operon1 | chr23_G1_Hap1:2965539-2971337 | G1 hap1 |
| | operon2 | chr23_G1_Hap1:3066486-3060688 | G1 hap1 |
| | operon3 | chr18_G1_Hap1:1244037-1238217 | G1 hap1 |
| | operon4 | chr18_G1_Hap1:1445816-1451637 | G1 hap1 |
| | operon5 | chr18_G1_Hap1:1557047-1551226 | G1 hap1 |
| | operon6 | chr18_G1_Hap1:1631553-1637381 | G1 hap1 |
| | operon7 | chr18_G1_Hap1:1877466-1871638 | G1 hap1 |

| | | | |
|---|---|---|---|
| | operon8 | chr17_G1_Hap1:1315313-1309495 | G1 hap1 |
| | operon9 | chr17_G1_Hap1:1343127-1337093 | G1 hap1 |
| | operon10 | chr2_G1_Hap1:5632794-5638615 | G1 hap1 |
| | operon11 | chr9_G1_Hap1:1019906-1014104 | G1 hap1 |
| 10 | operon1 | chr23_G1_Hap2:3348848-3354646 | G1 hap2 |
| | operon2 | chr23_G1_Hap2:3452200-3443996 | G1 hap2 |
| | operon3 | chr17_G1_Hap2:1316163-1310345 | G1 hap2 |
| | operon4 | chr18_G1_Hap2:1199255-1193435 | G1 hap2 |
| | operon5 | chr18_G1_Hap2:1401034-1406855 | G1 hap2 |
| | operon6 | chr18_G1_Hap2:1512240-1506419 | G1 hap2 |
| | operon7 | chr18_G1_Hap2:1586746-1592574 | G1 hap2 |
| | operon8 | chr18_G1_Hap2:1832659-1826831 | G1 hap2 |
| | operon9 | chr2_G1_Hap2:5622290-5628111 | G1 hap2 |
| | operon10 | chr9_G1_Hap2:1116432-1110629 | G1 hap2 |
| 9 | operon1 | chr23_SL1_Hap1:3643972-3649774 | SL1 hap1 |
| | operon2 | chr23_SL1_Hap1:3708666-3714468 | SL1 hap1 |
| | operon3 | chr18_SL1_Hap1:1548583-1554399 | SL1 hap1 |
| | operon4 | chr18_SL1_Hap1:1840053-1845869 | SL1 hap1 |
| | operon5 | chr18_SL1_Hap1:1962979-1968804 | SL1 hap1 |
| | operon6 | chr18_SL1_Hap1:1989172-1994997 | SL1 hap1 |
| | operon7 | chr17_SL1_Hap1:1766347-1772166 | SL1 hap1 |
| | operon8 | chr2_SL1_Hap1:5092921-5098743 | SL1 hap1 |
| | operon9 | chr9_SL1_hap1:1422798-1428601 | SL1 hap1 |

| | | | |
|---|---|---|---|
| 9 | operon1 | chr23_SL1_Hap2:3549336-3555137 | SL1 hap2 |
| | operon2 | chr23_SL1_Hap2:3628036-3622235 | SL1 hap2 |
| | operon3 | chr17_SL1_Hap2:1580482-1574663 | SL1 hap2 |
| | operon4 | chr18_SL1_Hap2:1735690-1741508 | SL1 hap2 |
| | operon5 | chr18_SL1_Hap2:1498935-1504755 | SL1 hap2 |
| | operon6 | chr18_SL1_Hap2:1807533-1813349 | SL1 hap2 |
| | operon7 | chr18_SL1_Hap2:1832243-1838068 | SL1 hap2 |
| | operon8 | chr2_SL1_Hap2:5378161-5383981 | SL1 hap2 |
| | operon9 | chr9_SL1_Hap2:1229757-1235560 | SL1 hap2 |

### Supplemental Table 2: Gene, repeat and secreted protein density in A and B compartments. Simple repeats were excluded.

| Isolate | Haplotype | Compartment | % gene sequence | % repetitive sequence | Genes per 10 Kb | | | |
|---|---|---|---|---|---|---|---|---|
| | | | | | All genes | Encoding secreted proteins | Encoding apoplastic effectors | Encoding cytoplasmic effectors |
| A4 | MAT-2 | A | **30.2%** | 45.7% | 2.2 | 0.07 | 0.01 | 0.04 |
| | | B | 16.1% | **59.9%** | 1.3 | 0.06 | 0.01 | 0.03 |
| | MAT-1 | A | **30.3%** | 45.9% | 2.2 | 0.07 | 0.01 | 0.04 |
| | | B | 16.4% | **59.8%** | 1.3 | 0.06 | 0.01 | 0.03 |
| A5 | MAT-6 | A | **29.6%** | 44.3% | 2.1 | 0.07 | 0.01 | 0.04 |
| | | B | 15.1% | **56.9%** | 1.3 | 0.06 | 0.009 | 0.03 |
| | MAT-3 | A | **29.5%** | 44.1% | 2.1 | 0.08 | 0.01 | 0.04 |
| | | B | 15.1% | **56.7%** | 1.3 | 0.06 | 0.007 | 0.03 |
| G1 | MAT-5 | A | **25.9%** | 48.2% | 1.9 | 0.07 | 0.01 | 0.04 |
| | | B | 19.1% | **55.2%** | 1.4 | 0.06 | 0.01 | 0.03 |
| | MAT-1 | A | **29.8%** | 45.6% | 2.1 | 0.07 | 0.01 | 0.04 |
| | | B | 17% | **56.5%** | 1.3 | 0.06 | 0.01 | 0.04 |
| SL1 | MAT-5 | A | **30.6%** | 43.8% | 2.2 | 0.08 | 0.009 | 0.04 |
| | | B | 17.9% | **58.9%** | 1.4 | 0.06 | 0.01 | 0.04 |
| | MAT-1 | A | **31.8%** | 41.7% | 2.3 | 0.08 | 0.01 | 0.04 |
| | | B | 18% | **59.8%** | 1.4 | 0.07 | 0.01 | 0.04 |

**Supplemental Table 3: The orthologous groups of effector fold proteins and the haplotype genes for A4, A5 and G1 heterokaryotic isolates.**

| A4 | | | |
|---|---|---|---|
| Orthogroup | Hap1 gene name | Hap2 gene name | MycFOLD |
| ORTHOMCL7270 | A4FUN_005280 | A4FUN_006146 | MycFOLD-17 |
| ORTHOMCL15332 | A4FUN_008862 | A4FUN_009580 | MycFOLD-16 |
| ORTHOMCL9391 | A4FUN_008861 | A4FUN_009569 | MycFOLD-15 |
| ORTHOMCL15175 | A4FUN_020662 | A4FUN_021189 | MycFOLD-14 |
| ORTHOMCL14639 | A4FUN_027365 | A4FUN_027926 | MycFOLD-12 |
| ORTHOMCL6090 | A4FUN_039991 | A4FUN_040843 | MycFOLD-11 |
| ORTHOMCL7824 | A4FUN_040107 | A4FUN_040966 | MycFOLD-10 |
| ORTHOMCL10045 | A4FUN_040113 | A4FUN_040971 | MycFOLD-9 |
| ORTHOMCL5763 | A4FUN_040233 | A4FUN_041081 | MycFOLD-8 |
| ORTHOMCL9535 | A4FUN_040125 | A4FUN_040983 | MycFOLD-7 |
| ORTHOMCL7477 | A4FUN_040115 | A4FUN_040972 | MycFOLD-6 |
| ORTHOMCL5451 | A4FUN_040124 | A4FUN_040982 | MycFOLD-5 |
| ORTHOMCL9535 | A4FUN_040125 | A4FUN_040983 | MycFOLD-4 |
| ORTHOMCL4093 | A4FUN_040126 | A4FUN_040984 | MycFOLD-3 |
| ORTHOMCL6128 | A4FUN_027326 | A4FUN_027902 | MycFOLD-2 |
| **A5** | | | |
| ORTHOMCL9814 | A5FUN_004912 | A5FUN_005690 | MycFOLD-17 |
| ORTHOMCL5094 | A5FUN_008000 | A5FUN_008564 | MycFOLD-16 |
| ORTHOMCL9938 | A5FUN_008010 | A5FUN_008565 | MycFOLD-15 |
| ORTHOMCL2740 | A5FUN_018569 | A5FUN_019077 | MycFOLD-14 |
| ORTHOMCL9429 | A5FUN_023685 | A5FUN_024229 | MycFOLD-13 |
| ORTHOMCL10401 | A5FUN_024523 | A5FUN_025050 | MycFOLD-12 |
| ORTHOMCL1678 | A5FUN_035706 | A5FUN_036440 | MycFOLD-11 |
| ORTHOMCL9429 | A5FUN_023685 | A5FUN_024229 | MycFOLD-1 |
| **G1** | | | |
| ORTHOMCL7888 | G1FUN_004776 | G1FUN_005695 | MycFOLD-17 |
| ORTHOMCL1302 | G1FUN_008110 | G1FUN_008678 | MycFOLD-16 |
| ORTHOMCL10457 | G1FUN_008111 | G1FUN_008679 | MycFOLD-15 |
| ORTHOMCL11404 | G1FUN_010370 | G1FUN_011142 | MycFOLD-14 |
| ORTHOMCL14483 | G1FUN_023906 | G1FUN_024548 | MycFOLD-13 |
| ORTHOMCL12498 | G1FUN_024871 | G1FUN_025377 | MycFOLD-12 |
| ORTHOMCL1850 | G1FUN_036183 | G1FUN_036862 | MycFOLD-11 |
| ORTHOMCL2742 | G1FUN_036284 | G1FUN_036969 | MycFOLD-10 |
| ORTHOMCL14483 | G1FUN_023906 | G1FUN_024548 | MycFOLD-3 |

**Supplemental Table 4**: **List of Go-Terms represented in each column shown in Figure 4C.**

**(Top. Biological Regulation; Bottom, Multicellular organismal process)**

| |
|---|
| biological regulation |
| regulation of biological process |
| multi-organism process |
| negative regulation of biological process |
| response to stimulus |
| cellular process |
| signaling |
| reproductive process |
| reproduction |
| metabolic process |
| positive regulation of biological process |
| localization |
| cellular component organization or biogenesis |
| developmental process |
| multicellular organismal process |

**Supplemental Table 5.** Root length colonization of *L. japonicus* with the four *R. irregularis* strains

| Strain | Sample | Total colonization | Internal Hyphae | Arbuscules | Vesicles |
|---|---|---|---|---|---|
| A5 | 1 | 94% | 94% | 66% | 70% |
| | 2 | 79% | 79% | 65% | 67% |
| | 3 | 94% | 94% | 87% | 80% |
| | 4 | 90% | 90% | 85% | 69% |
| A4 | 1 | 100% | 100% | 94% | 94% |
| | 2 | 99% | 99% | 98% | 83% |
| | 3 | 100% | 100% | 100% | 92% |
| | 4 | 93% | 93% | 89% | 78% |
| G1 | 1 | 97% | 97% | 93% | 62% |
| | 2 | 86% | 86% | 83% | 63% |
| | 3 | 99% | 99% | 96% | 79% |
| | 4 | 87% | 87% | 84% | 68% |
| SL1 | 1 | 52% | 52% | 49% | 37% |
| | 2 | 52% | 52% | 51% | 44% |
| | 3 | 80% | 80% | 80% | 72% |
| | 4 | 87% | 87% | 82% | 68% |

**Supplemental Table 6.** Root length colonization of *B. distachyon* with the four *R. irregularis* strains

| Strain | Sample | Total colonization | Internal Hyphae | Arbuscules | Vesicles |
|---|---|---|---|---|---|
| A5 | 1 | 66% | 66% | 44% | 24% |
| | 2 | 69% | 69% | 59% | 25% |
| | 3 | 62% | 62% | 45% | 28% |
| | 4 | 60% | 60% | 38% | 23% |
| A4 | 1 | 74% | 74% | 59% | 39% |
| | 2 | 83% | 83% | 59% | 48% |
| | 3 | 73% | 73% | 67% | 33% |
| | 4 | 82% | 82% | 68% | 49% |
| G1 | 1 | 57% | 57% | 43% | 20% |
| | 2 | 56% | 56% | 39% | 27% |
| | 3 | 66% | 66% | 47% | 29% |
| | 4 | 60% | 60% | 53% | 13% |
| SL1 | 1 | 51% | 51% | 46% | 10% |
| | 2 | 57% | 57% | 42% | 16% |
| | 3 | 67% | 67% | 55% | 16% |
| | 4 | 62% | 62% | 44% | 24% |
